# Applying knowledge-driven mechanistic inference to toxicogenomics

**DOI:** 10.1101/782011

**Authors:** Ignacio J. Tripodi, Tiffany J. Callahan, Jessica T. Westfall, Nayland S. Meitzer, Robin D. Dowell, Lawrence E. Hunter

## Abstract

Government regulators and others concerned about toxic chemicals in the environment hold that a mechanistic, causal explanation of toxicity is strongly preferred over a statistical or machine learning-based prediction by itself. Elucidating a mechanism of toxicity is, however, a costly and time-consuming process that requires the participation of specialists from a variety of fields, often relying on animal models. We present an innovative mechanistic inference framework (MechSpy), which can be used as a hypothesis generation aid to narrow the scope of mechanistic toxicology analysis. MechSpy generates hypotheses of the most likely mechanisms of toxicity, by combining a semantically-interconnected knowledge representation of human biology, toxicology and biochemistry with gene expression time series on human tissue. Using vector representations of biological entities, MechSpy seeks enrichment in a manually-curated list of high-level mechanisms of toxicity, represented as biochemically- and causally-linked ontology concepts. Besides predicting the canonical mechanism of toxicity for many well-studied compounds, we experimentally validated some of our predictions for other chemicals without an established mechanism of toxicity. This framework can be modified to include additional mechanisms of toxicity, and is generalizable to other types of mechanisms of human biology.

**Author summary:** Several recent computational methods have displayed excellent performance in predicting toxicity outcomes [1–3] of chemicals. Yet, to our knowledge, there is to date no computational approach to generate mechanistic hypotheses to answer **why** these chemicals elicit a toxic response. There is great value in understanding the mechanism of toxicity for a chemical that appears to elicit an adverse response. Novel small molecule development is one example, where a chemical that failed initial toxicological screenings could be assessed to evaluate the actual mechanism of toxicity, greatly reducing research time and expenses on subsequent ones. The value of a mechanistic awareness of toxicity also applies to pharmacovigilance, when researching rare adverse effects of a drug in subsets of the population. The development of oncological chemotherapeutics is another example, where certain mechanisms of cytotoxicity can actually be desirable to eliminate different types of tumor cells. More importantly, the costs, time expenditure, and ethical concerns of toxicity animal models, make in vitro and in silico approaches an enticing alternative. We present a solution that uses a combination of gene expression assays and biomedical knowledge to address the gap of answering the **why** question.

## Introduction

We currently have access to a wealth of biological, chemical and medical structured information represented in domain-specific ontologies. These ontologies elucidate diverse relationships between its constituent entities, like protein-protein interactions, or participation of enzymes (or chemicals) in biological processes, at specific cellular compartments. The Gene Ontology [4, 5] (GO) is the most commonly used, and a variety of tools exist to seek enrichment of particular concepts [6–8] or even specific pathways [9]. Our knowledge of molecular biology and biochemistry, however, goes well beyond what is described in GO, and it can be complemented by other ontologies and public databases that describe interactions between enzymes and/or chemicals.

In the case of in vitro toxicology studies involving gene expression assays (also referred to as “transcriptomics”), the simplest approach is to take the results of differential transcriptomics analysis and seek GO concept enrichment to gain clues of the underlying biological behavior of the treated cells. This is sometimes useful, yet often provides a somewhat disconnected list of terms that vary in specificity. The GO enrichment approach could be expanded by seeking enrichment of particular pathways, among databases like the Kyoto Encyclopedia of Genes and Genomes [10] (KEGG) or Reactome [11]. However, this output results in a list of possibly related biological pathways that are not necessarily tailored to the study’s domain (e.g. toxicology). Moreover, current pathway enrichment strategies don’t take into account the sequential order in which the experimentally-significant expression changes occur.

The use of artificial intelligence to infer mechanistic behavior in biology or other disciplines is still at a nascent stage. Most of the computational work on biological mechanisms has been focused on their representation [12, 13], and studies aimed at elucidating a mechanism of toxicity have generally been targeted to specific compounds, or narrow classes of them. Prior work in seeking enrichment of adverse outcome pathways (AOPs [14], which are the most common representation for chemical-specific toxicity), was focused on targets like pulmonary fibrosis [15] or fatty liver [16]. Computational approaches are sometimes faced with skepticism, particularly as the techniques become increasingly “black box”-like [17]. A hypothesis generation tool that produces mechanistic narratives backed by curated existing knowledge, however, would result in a more attractive alternative as scientists can validate the plausible explanations offered. Moreover, scientists could highlight any potential mistakes, adjusting our web of knowledge accordingly, and having the entire community benefit from it.

The idea of computationally-generated explanations is not new, as Schank [18] proposed a framework for this in 1986. At that time, the breadth of scope and lack of computational power resulted in explanation patterns remaining mostly as a theoretical exercise. However, we now have the tools to make it possible. Here we present a computational framework (Fig. 1) to produce mechanistic hypotheses of toxicity from in vitro assays. Taking a time series gene expression experiment as input, MechSpy uses a graph representation of our current knowledge of molecular biology and biochemistry, to generate a transparent narrative for each of the three most likely mechanisms of toxicity taking place, linking experimental events to different mechanism steps.

**Fig 1.**
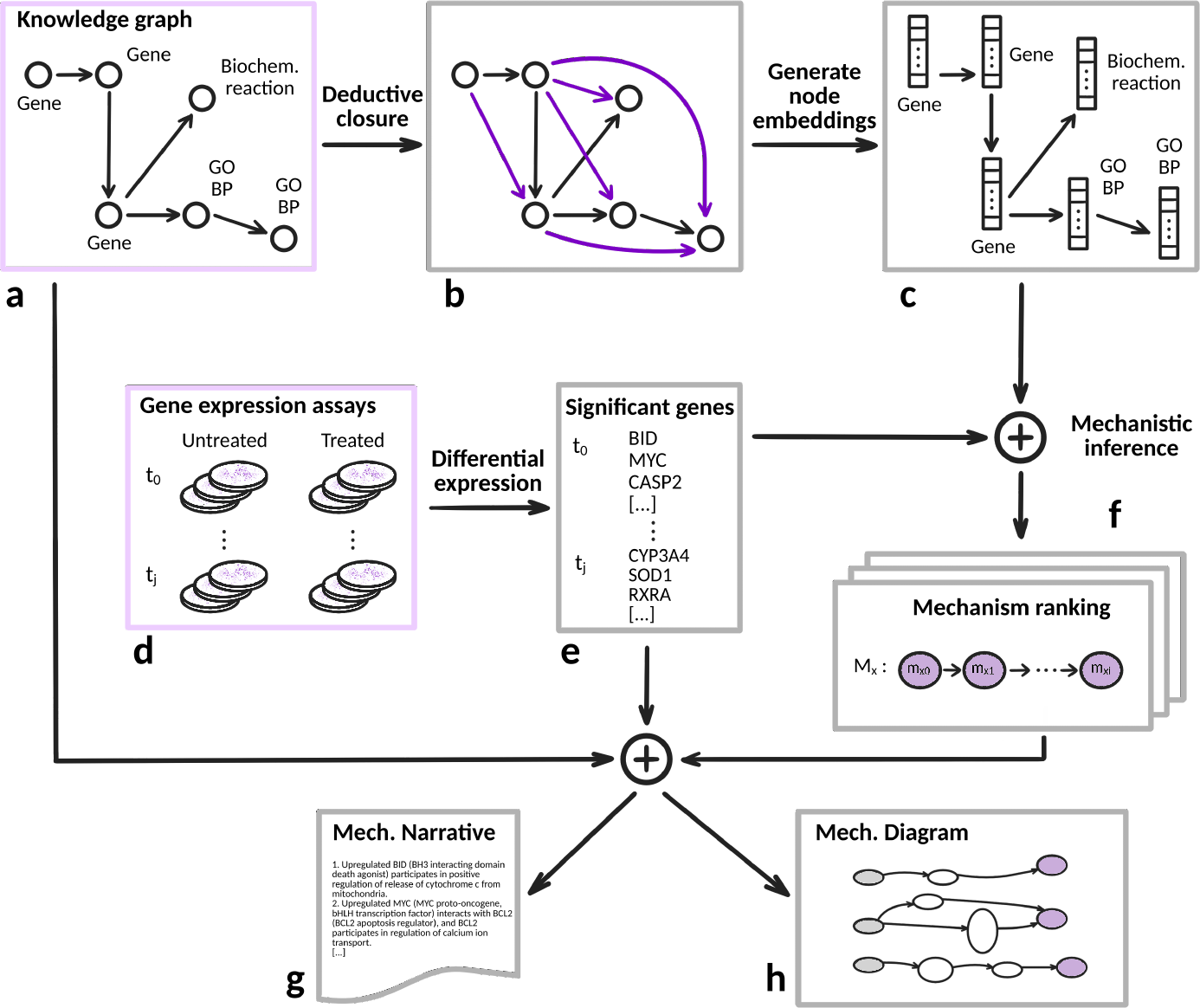
Overview of MechSpy’s mechanistic inference process. The knowledge graph (**a**) of semantically-integrated ontologies and databases and the transcriptomics data (**d**), in light purple frames, are our inputs. After adding new edges to the graph by deductively closing it (**b**), MechSpy uses node2vec to generate dense vector embeddings of each node (**c**). We also perform differential expression analysis on the transcriptomics data (**d**), and obtain a list of the top N most significant changes in gene expression (**e**). Based on these changes, and using the embeddings for genes and all mechanism steps, MechSpy generates an enrichment score for each mechanism (**f**). Using the original knowledge graph (**a**) and the significant genes across time (**e**), it then produces both a narrative (**g**) and a graphical explanation (**h**) for each of the three most enriched mechanism.

## Materials and methods

### Toxicity mechanisms

Many mechanisms in biology can be represented as an ordered sequence of ontology concepts. After an extensive literature review, we curated a list of high-level mechanistic toxicology descriptions. A total of 11 high-level mechanisms were curated, based on a mechanistic toxicology textbook [19], which we were able to represent as causally-linked concepts from the gene ontology. After parsing the textual description of a mechanism, we listed the sequence of key events that would summarize it at a high level. For each of these events, we looked for the GO concept that most closely represented it. The mechanisms with their respective steps, which represent events at a molecular level that must follow a sequential order, are listed in Table. 1. We attempted to capture with these some of the most common ways a toxicological insult results in an adverse cellular outcome.

**Table 1.**
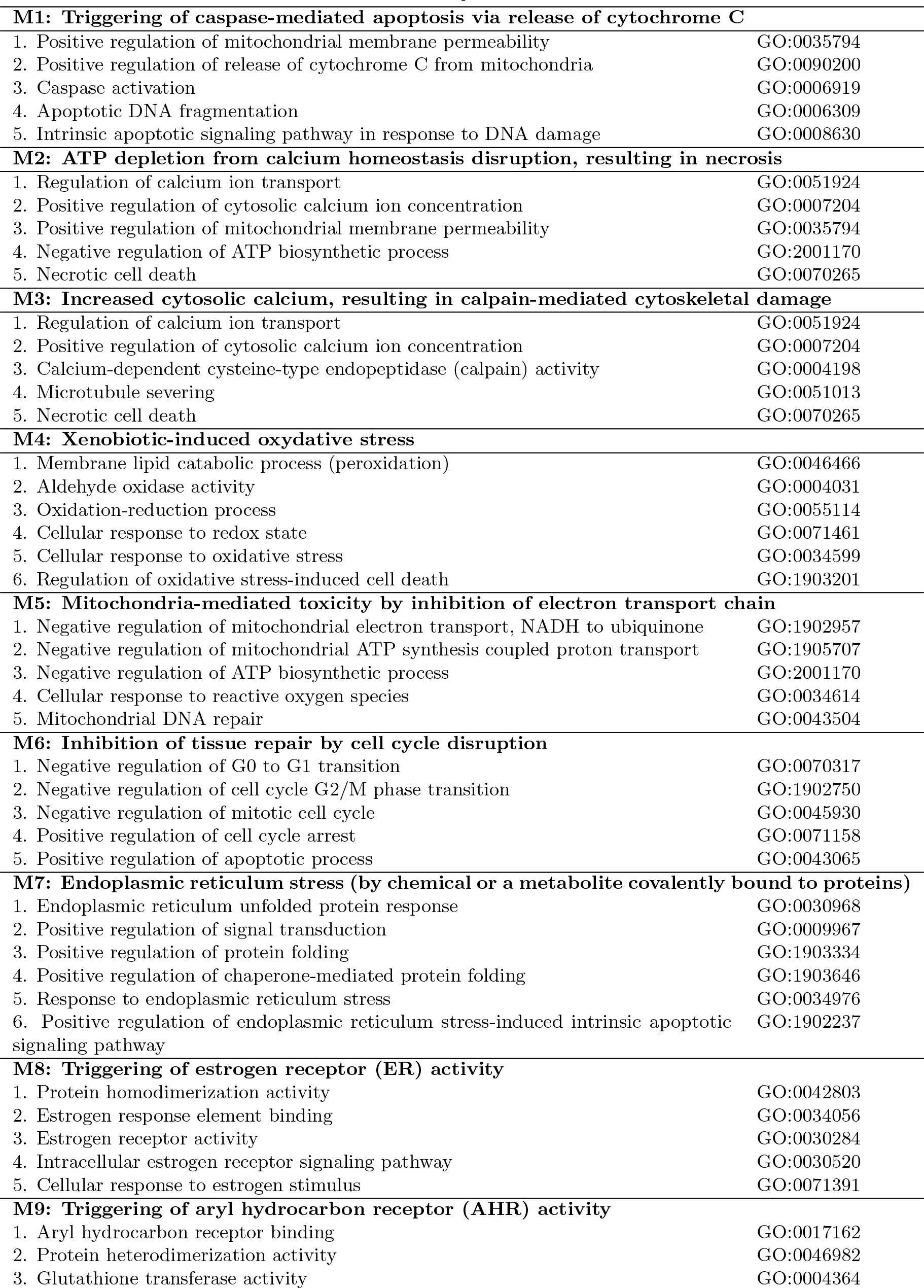
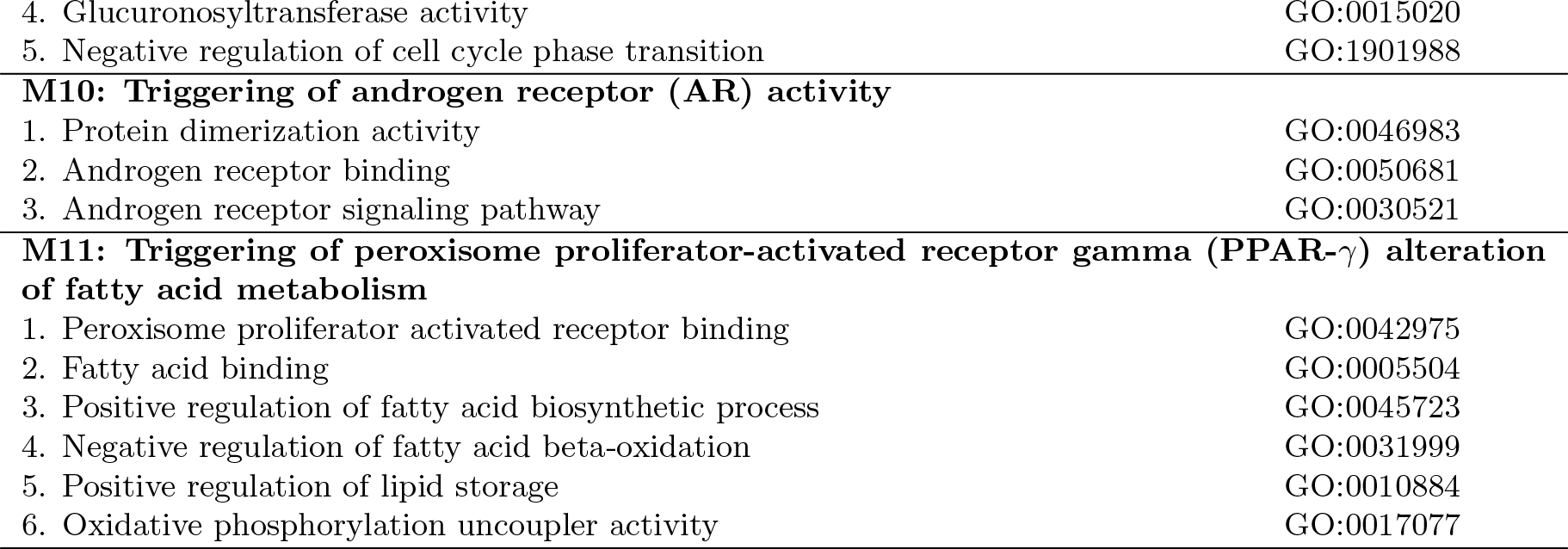
Mechanisms of toxicity evaluated. These mechanisms of toxicity were manually curated from the literature, and every mechanism step was represented using an ontology concept. The enrichment of each mechanism step is expected to happen following the sequential order in which they are described here.

These mechanisms can be considered as cell-focused, different from AOPs which go beyond the cellular scope and towards organ- and individual-specific responses. These mechanisms were reviewed by several members of the Toxicology department at the University of Colorado, Anschutz medical campus. We surveyed the literature for evidence of mechanistic explanations of toxicity for every compound used in the exposure assays we tested. The possible mechanism labels and literature sources for each of these chemicals are listed in Supplemental Table S1. MechSpy selects the three most likely high-level mechanisms of toxicity for each transcriptomics time series, and produces a putative explanation for each.

### Knowledge graph

A knowledge graph (KG) is a more powerful tool to employ than any ontology by itself, as it provides much richer contextual information for any concept, and can uncover relations between entities that would be missed in separate ontologies. We extended a KG [20] (Fig. 1.a) that was generated by semantically integrating multiple open biomedical ontologies (OBOs [21]) and other sources of publicly available linked open data. An ontology is a formal representation of entities or concepts, and the relations between them. Ontologies employ a directed acyclic graph (DAG) representation, usually described as a list of “triples” (subject, predicate, object). The KG included concepts from the Gene Ontology [4, 5], Protein Ontology [22], Cell Ontology [23], Human Phenotype Ontology [24], Human Disease Ontology [25], and Chemical Entities of Biological Interest [26], among others, in a semantically-consistent fashion. Only human entities and the relations between them were included. Several public databases were also used to incorporate additional directed edges into the KG, e.g. the Cellular Toxicogenomics Database [27], Reactome [28], the STRING database [29], the AOP Wiki [30], the National Cancer Institute thesaurus [31], and Uniprot [32].

This KG focuses on concepts directly related to human biology, rather than those involving model organisms. (Fig. 2) illustrates the richness of information that can be extracted from these interconnected sources. Once the knowledge graph was built, the ELK reasoner [33, 34] was run to deductively close and complete the graph (Fig. 1.b), adding new edges for transitive relations where applicable. For example, the “DFFA” gene participates in the “Apoptosis-induced DNA fragmentation” pathway, and the “Apoptosis-induced DNA fragmentation” pathway has part in the “Apoptotic DNA fragmentation” biological process, so a new edge is added to represent “DFFA participates in the apoptotic DNA fragmentation biological process”. The graph was used both in its original form to generate the mechanistic narratives, and in its deductively-closed form to generate vector embeddings for its nodes.

**Fig 2.**
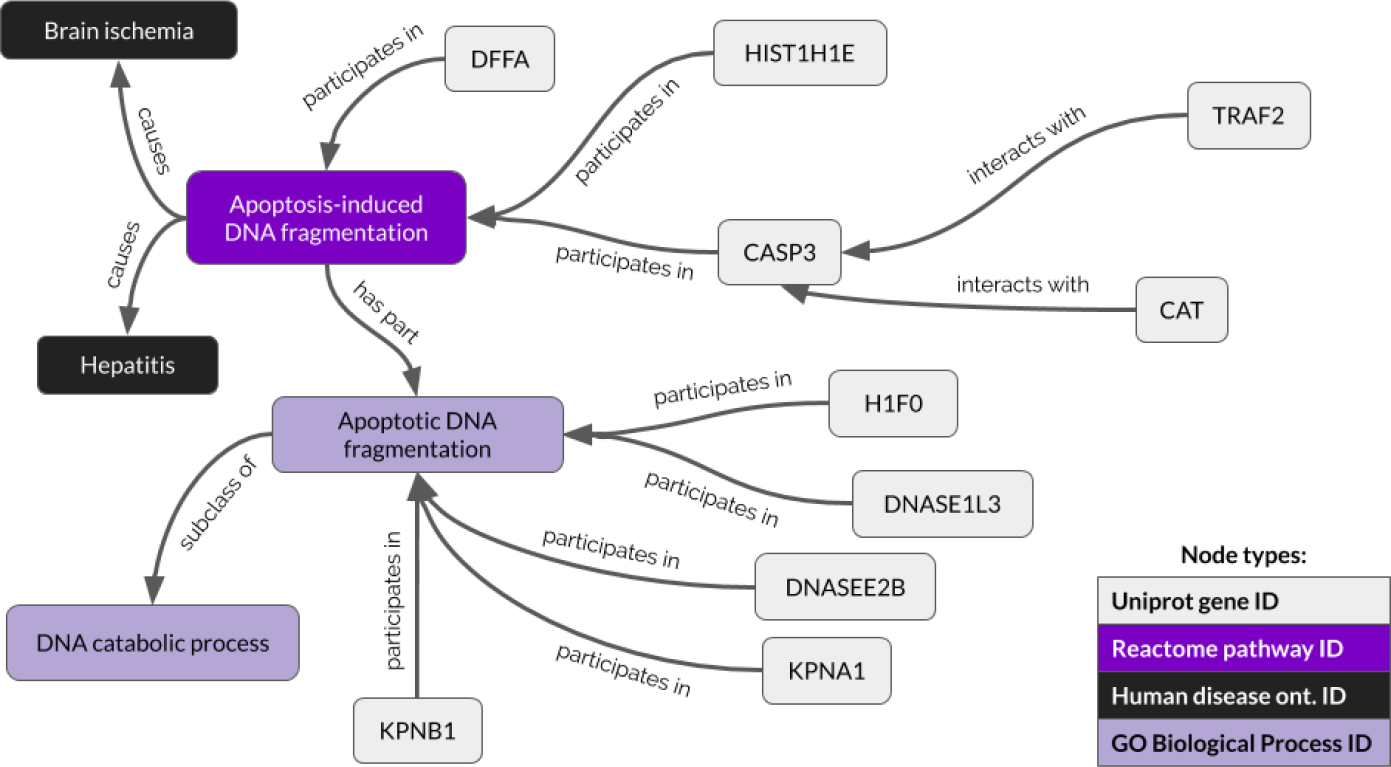
Illustrative knowledge graph sample. Example of how ontology concepts and database sources are interconnected in the KG extended from the PheKnowLator project [20].

### Data sources

Since the KG is focused on human biology, we sought out publicly available time series datasets that used human cells, from different tissue types. The raw human microarray data (Fig. 1.d) used as input was sourced from the Open TG-Gates [35] database, the carcinoGENOMICS [36] project and Netherlands Toxicogenomics Centre projects available in the diXa Data Warehouse [37], as well as other publicly available tobacco-related human microarray datasets [38–41] in ArrayExpress [42] that used human nasal, buccal and bronchial epithelial tissue. These sources provided a variety of time series gene expression experiments, at different time points and using different human cell types. Specifically, we used samples from Open TG-Gates that featured 3 time points, 3 replicates for each condition and 3 doses of exposure (low, medium, and high, varying by each chemical). Other samples from Open TG-Gates were also chosen as canonical examples of toxicity mechanisms from Urs A. Boelsterli’s textbook [19]. For all other data sources, we used any time series which had at least two time points with significant differentially-expressed genes.

We used gene expression time series from diXa studies DIXA-002 (liver cells HepaRG and HepG2), DIXA-003 (kidney cells RPTEC/TERT1), DIXA-004 (lung epithelial cells BR200, BR234, BR259, BR234/CDK4, and BR234/p16), and DIXA-078 (HepG2 liver cells). Using the Limma [43] R package with robust multiarray average (RMA) background correction and normalization, the control and treatment replicates were contrasted to determine the most significant gene expression changes at each time point, based on a p-value cutoff of 0.05 (Fig. 1.e). The only exception was for the DIXA-004 lung epithelium samples, since only a single replicate per condition was available, so we had to rely on fold change to determine the most significant genes. We evaluated a total of 239 different exposure time series, out of which 221 resulted in the prediction of at least one mechanism of toxicity with sufficient statistical confidence.

### Mechanistic inference

The inference step (Fig. 1.f) consists of scoring each curated mechanism, based on how much the affected gene nodes in the KG (corresponding to the differential analysis findings) relate to the GO nodes describing the mechanism steps. The three mechanisms of toxicity with the highest ranking are then presented as the most likely candidates. We used a latent vector representation (embedding) of each node in the KG, a popular tool in natural language processing applications [44], to determine mechanism enrichment scores. The idea of generating dense real number vector representations of language tokens can be applied to KGs, with techniques to generate embeddings for either nodes or edges. These are generated by models that predict the most likely node, based on the context (neighboring) nodes. Thus, two nodes with a similar vector representation in semantic space are expected to have a closely related meaning. A recent review [45] provides a good summary of the current models. One such algorithm to generate node embeddings from a graph is node2vec [46], which utilizes random walks from the node in question up to a number of hops away, with a bias for depth or breadth configurable by hyperparameters. This was our method of choice to create vector embeddings to represent the different kinds of entities represented in our graph (biological processes, genes, proteins, biochemical reactions, etc). Using an emphasis on breadth (`--directed --dimensions 32 --q 3`, for best performance with our deductively closed graph) we created 32-dimensional vector representations of all 109,255 nodes (Fig. 1.c).

Cosine distance is an established way to compare vector embeddings, and an appropriate metric for this study, since we focused on similar hyperdimensional orientation to denote similar meaning. For each experimental time point *j*, MechSpy collected the embeddings for the (up to) 100 most significant genes. These embeddings were then averaged to obtain the centroid 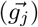 of all those gene vectors, which represented the significant expression changes, as a whole, at that time point (Fig 3). From this average vector 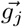, MechSpy calculated the cosine distance to each mechanism step of every curated mechanism. In order to use this as a weight rather than a distance (such that a larger magnitude equals a stronger enrichment), it substracted the cosine distance value from 1 to use it as an enrichment score (also known as cosine similarity).

**Fig 3.**
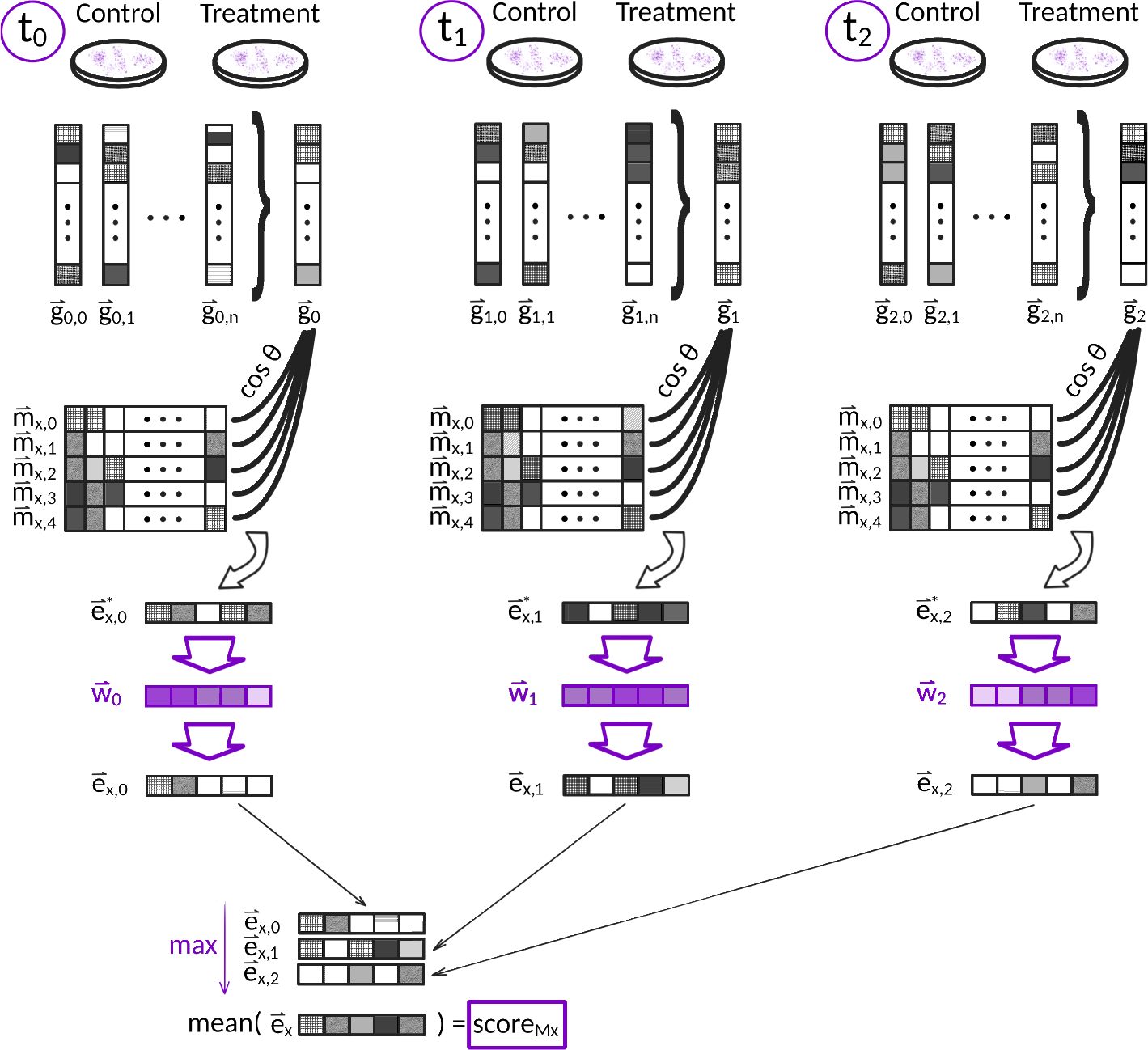
Summary of the mechanistic inference architecture, showing the use of vector embeddings generated using node2vec, and our sequential order penalty scheme to find the score of a particular mechanism. In this hypothetical example we have 3 experimental time points, and a mechanism composed of 5 causally-linked steps. For every time point, the *n* most significant gene changes (where 1 ≤ *n* ≤ 100) are averaged into a single vector. MechSpy then generates a preliminary enrichment vector 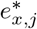, which consists of the cosine similarity value between that time point’s gene aggregation and each mechanism step, subtracted from 1. The sequential penalty filter (in purple, dividing the 5 mechanism steps in 3 bins) gives less weight to mechanism steps that don’t correspond to the time point in question, with an increasing penalty the farther away we are from our corresponding bin. Finally, the weighted enrichment vectors for each time point are combined such that the maximum score for each mechanism step is kept 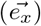, and the score for this mechanism is the mean of these maximum values.

The sequential order in which the mechanism steps are described (and enriched) matters. If a set of genes are closely related to the last step of a mechanism, for example, we value their contribution in the last time points of our experimental time series, more than in the first ones. Therefore, we devised a weighting scheme that prioritized the contributions to each mechanism step corresponding to the current time point (Fig. 3), in purple). Given a mechanism *M*_*x*_ = [*m*_*x*0_, *m*_*x*1_, …, *m*_*xi*_] with *i* steps, and an experimental time series *T* = [*t*_0_, *t*_1_,…, *t*_*j*_] with *j* time points, MechSpy segments *M* in |*T*| bins. These bins were used for a sequential weighting scheme, to penalize genes that enrich mechanism steps out of proper sequential order. A vector 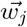 of size |*M*_*x*_| was used to weight enrichment scores at each time point. Having *b*_*j*_ as the bin index corresponding to the current time point, and *b*_*s*_ as the bin index being evaluated, the weight for each bin was calculated as:

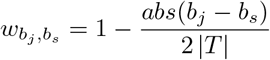

Thus, each of the values *e*_*xi,j*_ of an enrichment vector 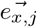 of |*M*_*x*_| dimensions (Fig.3, bottom), containing the enrichment score for each of the *i* steps in mechanism *x* at time point *j*, were calculated as:

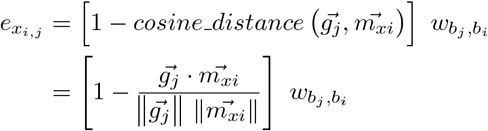

To test whether sequential order of enrichment improved our predictions, we pseudo-randomly shuffled time points for all time series (ensuring they were always in incorrect order), and recalculated all mechanism enrichment scores. The resulting accuracy was lower across all data stratifications, particularly for the top-scoring mechanism, which demonstrated the need for MechSpy’s sequential penalty scheme.

Since the cosine similarity and weight penalty are bounded in [0, 1], so is the enrichment score for each mechanism step. The maximum values of the enrichment vectors for each experimental time point *j* were then combined into a single overall enrichment vector 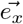 for mechanism *M*_*x*_ (Fig. 3, bottom). The 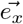 vector featured the highest scores achieved for each mechanism step. The final enrichment score for mechanism 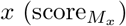 was the average of 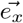 elements. The mechanisms used in this study were relatively short; if significantly longer mechanisms are defined, it would be advisable to calculate the median of all final mechanism step scores (rather than the mean) to obtain the final enrichment score, to prevent outliers from effecting a significant change. This procedure was repeated for every curated mechanism. Thus, our set of predictions *P* is defined as:

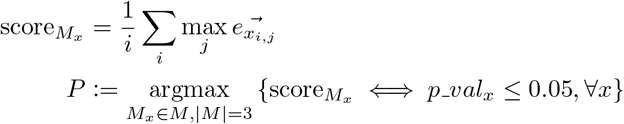

The three mechanisms with the highest score in *P* that were deemed statistically significant (see below) became the ones suggested as most likely.

### Statistical significance

In order to determine the statistical significance of these predictions, we evaluated how they would compare to a pseudo-random assortment of genes. To this end, we employed a bootstrap approach where MechSpy calculated the final score for each mechanism, using random draws of the same number of genes originally used at each time point. Given that all microarray public samples used the same chip (Affymetrix Human Genome U133 Plus 2.0), the genes were randomly drawn from all probes available in it, represented by their ontology concepts in the KG. After 1000 simulations, MechSpy generated an empirical distribution of mechanism scores under the equivalent experimental conditions, whose median it used to compare our real mechanism score against. The resulting p-value for each mechanistic prediction was therefore calculated as:

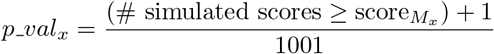

This empirical p-value, which at 1000 iterations has a lower bound of 9.99×10^−4^, was used to discard mechanistic predictions that were not significant, based on a cutoff of 0.05. Mechanisms that scored below the median of random simulations were also dropped. The rest of the mechanisms were sorted in descending order by their final enrichment score to determine the three most likely predictions.

### Performance evaluation

Chemicals can elicit different mechanisms of toxicity which can depend on dosage and tissue type. For nearly half of the assays evaluated, we have more than one possible correct label of expected mechanisms. We thus decided to present the top-three most likely mechanisms, considering that MechSpy is a hypothesis generation aid. The existence of multiple correct labels in addition to the lack of a real gold standard, made the evaluation difficult in traditional terms of performance and recall. To calculate precision, a “positive” sample was defined as having at least one of the expected mechanisms among the predictions evaluated (top, top-two or top-three).

We used the precision for all assays of the top-scoring mechanism, then among the top-two, and among the top-three, as a global performance metric. This metric was applicable to any experiment for which we had at least one statistically significant prediction. In other words, those assays for which the gene expression data was not sufficient to produce at least one mechanistic prediction, were ignored. The datasets were then stratified by two criteria: chemicals with only one known mechanism of toxicity, and chemicals with two known mechanisms of toxicity. Taking into account that not all assays we tested were conducted at toxic doses, these precision values represent a lower bound estimate of performance. Therefore, a third stratification dimension used was to consider only assays for the highest dose available, for each chemical and cell type combination. The results at high exposure doses likely reflect the most realistic performance of MechSpy.

### Mechanistic narratives

Another aspect of MechSpy’s novelty is that it’s not simply a mechanism prediction tool: for each toxicity mechanism, MechSpy produces a narrative (Fig. 1.g) of a putative explanation. Using the non-deductively-closed version of the KG, for each of the top-three ranked mechanisms, MechSpy searched for paths connecting the most significant genes at each time point *j* with each of the mechanism steps. It prioritized those up/downregulated genes at the time points that corresponded to each mechanism step (based on our binning strategy described in Methods). (Fig. 4) presents an example narrative generated for liver cells exposed to a 400*μ*M concentration dosage of diclofenac sodium for up to 24 hours, for which “ATP depletion due to calcium homeostasis disruption” (M2) was predicted as one of the most likely mechanism of toxicity.

**Fig 4.**
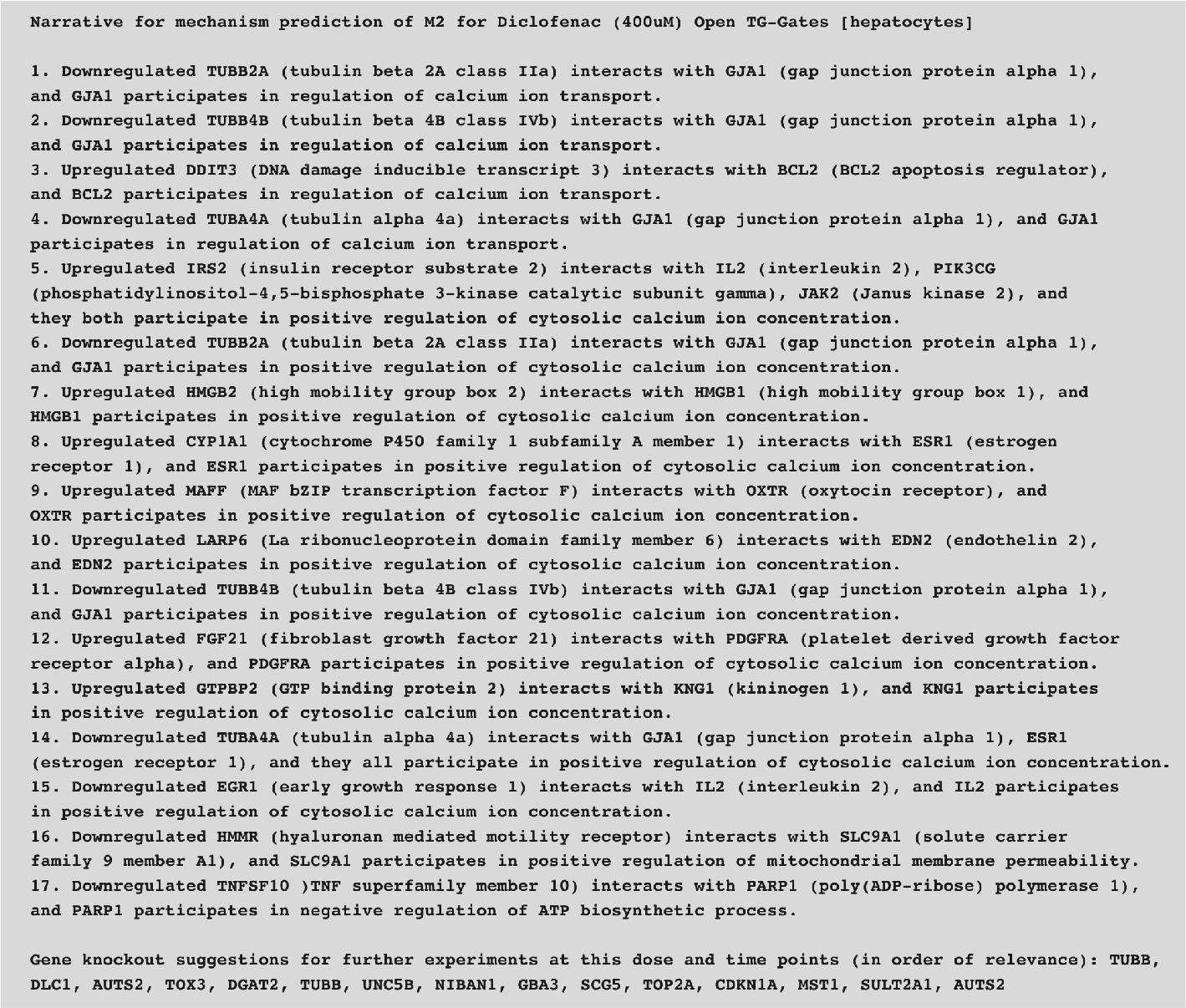
Mechanistic narrative generated for a time series of diclofenac sodium exposure. Example of a generated mechanistic narrative for a particular transcriptomics time series, generated by MechSpy.

This narrative was limited by default to paths from genes to mechanism steps no farther than two hops away in our KG (to focus on closely related entities), or more if intersecting at Reactome entities, such as pathways or biochemical reactions. This limitation can, however, be easily relaxed to include many other possible pathways. The mechanistic explanation also includes a list of suggested gene knockouts, to help experimentally validate these claims. This list is sorted by significance among all time points, and could help discover new genes with a key role in the enriched mechanism. A visualization of each putative mechanistic explanation (Fig. 1.h) is also presented as a network diagram. Fig 5 shows the generated diagram corresponding to the same example narrative presented in Fig. 4.

**Fig. 5.**
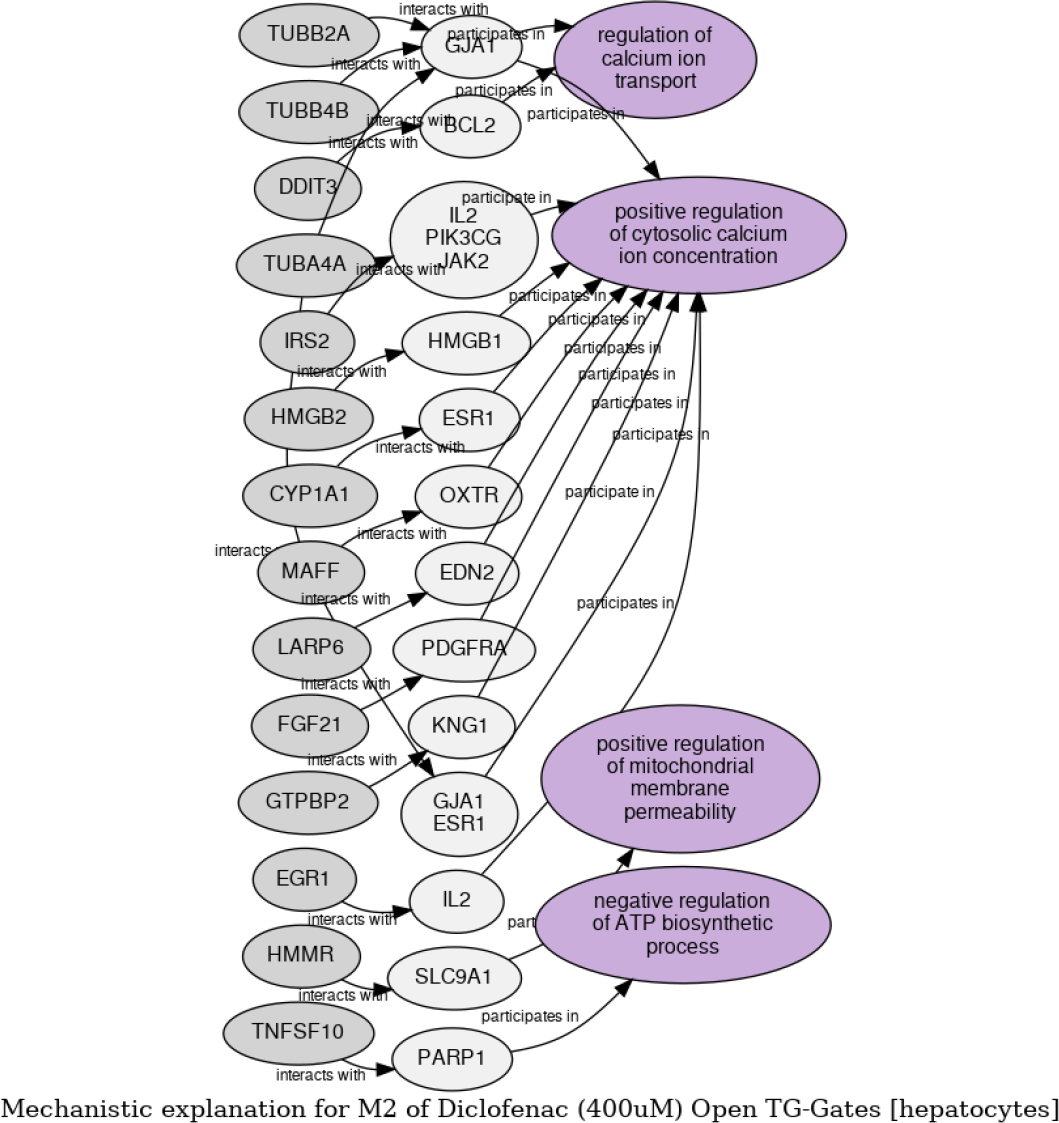
Graphical mechanistic explanation example, generated by MechSpy. The sequence of events should be followed top to bottom, left to right. Nodes in dark gray are the significant gene changes along the multiple time points (in order), and purple nodes represent the mechanism steps enriched.

### Experimental validation

We sought out experimental validation for two of the chemicals without a well-known mechanism of toxicity: adapin and chlorpromazine. The selection of chemicals and mechanistic hypotheses to test was based on mechanisms consistently predicted by MechSpy at all concentrations, or at least at the two higher concentrations on the tissue used in the time series. The MITO-ID kit from Enzo Life Sciences was used to assess mitochondrial-mediated toxicity. This kit measures a shift in mitochondrial membrane potential from excited membrane to depolarized membrane (both markers of mitochondrial toxicity), and compromised plasma membrane integrity (a marker of cell death). We validated our predictions of mitochondial toxicity using HUH7, a hepatocyte-derived carcinoma cell line. These results were also confirmed on an additional hepatocyte cell line, HepG2, and further evaluated on an orthogonal cell line (HCT116, human colorectal carcinoma cells) to assess tissue specificity. For experimental details and outcomes in all cell lines, see Supplemental Experimental Details.

## Results

From the time series assays (several hours apart) MechSpy has been able to predict the most likely mechanisms of toxicity, starting with a set of compounds for which a “canonical” mechanism has been established (based on a mechanistic toxicology textbook [19]). As a proof of concept, we utilized assays from Open TG-Gates [35] that used chemicals with an established most likely mechanism in the literature [19], and for which we had at least two time points with any significantly up/down-regulated genes. For this subset of experiments, we could predict the canonical mechanism(s) as the top choice with a precision of 0.594, and of 0.812 among both the top-two and top-three. This subset of experiments included exposure concentrations that were most likely not toxic. Looking only at the highest dose used for each compound, our precision for the most likely mechanism is 0.615 and we were guaranteed to make a correct prediction among the top-two most likely already. With these encouraging results, we moved on to include the rest of chemicals in Open TG-Gates for which we had 3 time points, 3 replicates and 3 exposure doses (low, medium and high, which varied substantially depending on the compound), as well as other public gene expression time series (diXa-002, diXa-003, diXa-004, diXa-078, E-MTAB-4740, E-MTAB-4742, E-MTAB-5157, and E-MTAB-5697).

For all experiments using chemicals which we could label with one or more possible mechanisms based on the scientific literature, one or more of our top-three predictions matched its expected label with a precision of 0.747. If we only considered the highest dose per chemical and cell type, this precision raised to 0.843. A sample of MechSpy’s predictions with their respective empirical p-values is shown in Table. 2. Besides looking at the highest dose for each chemical and cell type, we further stratified the assays regarding whether there was only one known mechanism of toxicity, or two at most. The performance for these various ways to segment the public datasets are displayed in Table. 3. Taking the sequential order of the mechanism steps in mind, when calculating the enrichment scores, contributed to this performance. The accuracy obtained was also significantly better than a baseline estimated from random draws of three mechanisms, for every compound with known mechanisms of toxicity (Fig. 6).

**Table 2.** MechSpy predictions for a subset of the time series evaluated.

| Chemical used | Dose | Known mechanisms | #1 Predicted mechanism | #2 Predicted mechanism | #3 Predicted mechanism |
| --- | --- | --- | --- | --- | --- |
| acetaminophen | 200μM | M2, M4, M5 | M1 (6.99E-03) | **M4** (8.99E-03) | N/A |
| acetaminophen | 1mM | M2, M4, M5 | **M4** (9.99E-04) | **M5** (9.99E-04) | M1 (9.99E-04) |
| acetaminophen | 5mM | M2, M4, M5 | **M4** (9.99E-04) | **M2** (9.99E-04) | M1 (9.99E-04) |
| aflatoxin B1 | 0.24μM | M4 | M5 (9.99E-04) | M1 (9.99E-04) | M2 (9.99E-04) |
| aflatoxin B1 | 1.2μM | M4 | **M4** (9.99E-04) | M5 (9.99E-04) | M1 (9.99E-04) |
| aflatoxin B1 | 6μM | M4 | **M4** (9.99E-04) | M5 (9.99E-04) | M1 (9.99E-04) |
| benzyl alcohol | 10mM | M1 | **M1** (9.99E-04) | M4 (9.99E-04) | M5 (9.99E-04) |
| cyclosporin A | 1.2μM | M1, M5 | **M5** (9.99E-04) | M4 (1.40E-02) | **M1** (9.99E-04) |
| cyclosporin A | 6μM | M1, M5 | **M5** (9.99E-04) | **M1** (9.99E-04) | M2 (3.00E-03) |
| diclofenac | 16μM | M2, M7 | **M2** (9.99E-04) | M4 (6.99E-03) | M8 (7.99E-03) |
| doxorubicin | 2μM | M5 | M4 (9.99E-04) | M1 (9.99E-04) | M2 (9.99E-04) |
| doxorubicin | 10μM | M5 | **M5** (9.99E-04) | M4 (9.99E-04) | M1 (9.99E-04) |
| glibenclamide | 4μM | M2 | M4 (9.99E-04) | M5 (3.00E-03) | M1 (3.00E-02) |
| glibenclamide | 20μM | M2 | M4 (9.99E-04) | **M2** (9.99E-04) | M5 (2.00E-03) |
| imipramine | 100μM | M1, M11 | M9 (7.99E-03) | M10 (2.00E-03) | **M11** (2.00E-02) |
| isoniazid | 400μM | M4 | **M4** (9.99E-04) | M2 (9.99E-04) | M1 (9.99E-04) |
| isoniazid | 2mM | M4 | **M4** (9.99E-04) | M2 (9.99E-04) | M5 (9.99E-04) |
| isoniazid | 10mM | M4 | **M4** (9.99E-04) | M2 (9.99E-04) | M5 (9.99E-04) |
| rotenone | 2μM | M5 | **M5** (9.99E-04) | M1 (9.99E-04) | M9 (9.99E-04) |
| sulfasalazine | 30μM | M4 | M5 (4.30E-02) | M2 (1.20E-02) | M3 (9.99E-04) |
| sulfasalazine | 150μM | M4 | **M4** (9.99E-04) | M5 (9.99E-04) | M2 (9.99E-04) |

Mechanistic inference results for a few of the 239 time series evaluated, that utilized chemicals with well-established mechanisms of toxicity at a variety of concentrations, some of which are examples in Boelsterli’s textbook [19]. The **bolded** predictions are those that match the known mechanisms for that chemical.

**Table 3.** MechSpy performance across multiple levels of stratification.

| Stratification | # time series | Top Precision | Top-2 Precision | Top-3 Precision |
| --- | --- | --- | --- | --- |
| Only one known mechanism | 117 | 0.376 | 0.598 | 0.632 |
| Only two known mechanisms | 88 | 0.432 | 0.648 | 0.852 |
| Highest dose per chemical/cell type | 108 | 0.463 | 0.694 | 0.843 |
| Highest dose per chemical/cell type, only one known mechanism | 61 | 0.443 | 0.705 | 0.754 |
| Highest dose per chemical/cell type, only two known mechanisms | 39 | 0.436 | 0.615 | 0.949 |
| All assays | 221 | 0.430 | 0.647 | 0.747 |

We display here the precision for the top-scoring mechanism, as well as among the top-two and top-three scoring mechanisms for time series.

**Fig 6.**
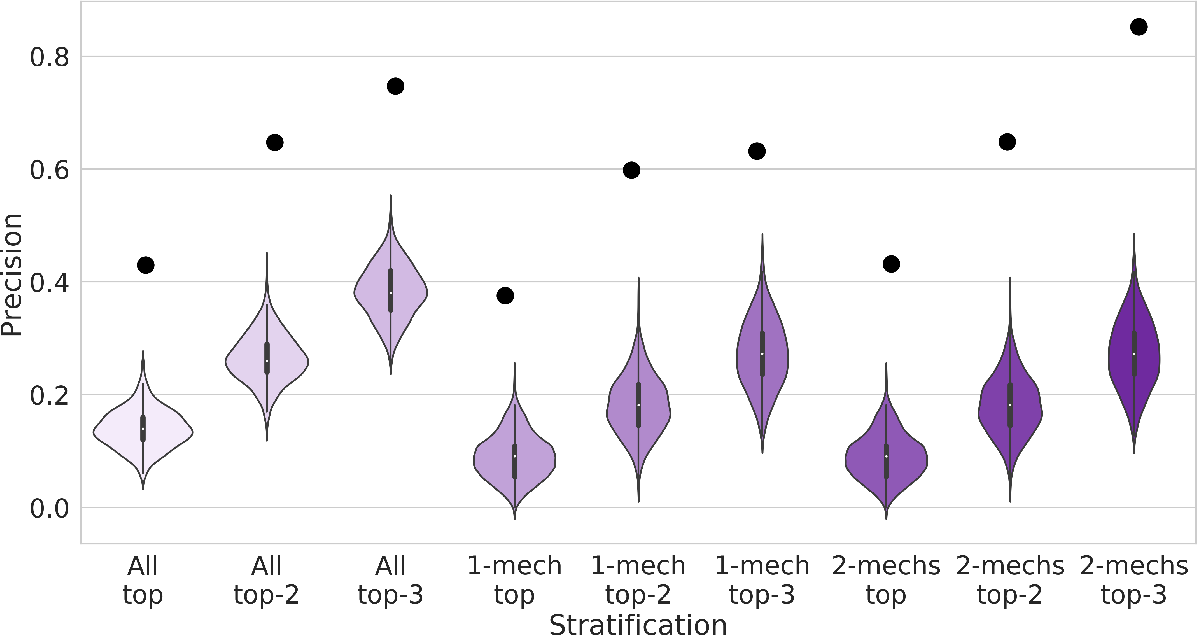
Simulations of baseline precision. Comparison of actual precision values (black dots, top) to baseline estimations from random mechanism draws (violin plots, bottom), for different segmentations of the data (see Table 3). For each chemical used in the public datasets with one or more known mechanisms of toxicity, we randomly drew three mechanisms of the eleven curated (without replacement) to simulate the top-three enrichments. The accuracy across all chemicals was then calculated, and the process was repeated 1000 times. The violin plots show the distribution of baseline accuracy scores from those 1000 runs.

It is worth reiterating that many of these chemicals don’t act via a single mechanism of toxicity, so the accuracy for the strongest enrichment score is actually a lower-bound estimate. The actual mechanistic landscape is likely better represented by a combination of these top enriched mechanisms. The full list of mechanistic hypotheses for all time series is available in Supplemental Table S2. Some of the predictions can be linked to known toxicity endpoint organs. Clonidine, for example, has known cardiotoxicity issues [47] at high doses. This is very common with the predicted mitochondria-mediated toxicity, since cardiomyocytes are highly energy-demanding cells and this adverse mechanism results in a sharp decrease in ATP synthesis. While the strongest hypothesis is that it triggers apoptosis via caspase release [48] (M1), MechSpy predicted it acted via mitochondrial-mediated toxicity (M5) with a higher score than M1 for all concentrations and cell types.

Some of the chemicals used in the public datasets don’t have a well-established mechanism of toxicity to date. We experimentally validated one of the generated mechanistic hypotheses for two of those, adapin and chlorpromazine. MechSpy consistently predicted mitochondrial-mediated toxicity for the two highest concentrations, as one of the three most likely mechanisms. Additionally, we exposed these chemicals to a human colorectal carcinoma cell line (HCT116), to evaluate whether the toxicity was specific to the cell types in question. MechSpy’s prediction that these chemicals affect liver tissue was confirmed using HUH7 cells and HepG2 hepatocyte-derived carcinoma cells, and tested on an unrelated cell line (HCT116) to assess tissue specificity. (Fig. 7) shows validation of our predictions using Huh7 cells, and the rest of assay outcomes are made available in Supplemental Experimental Details.

**Fig 7.**
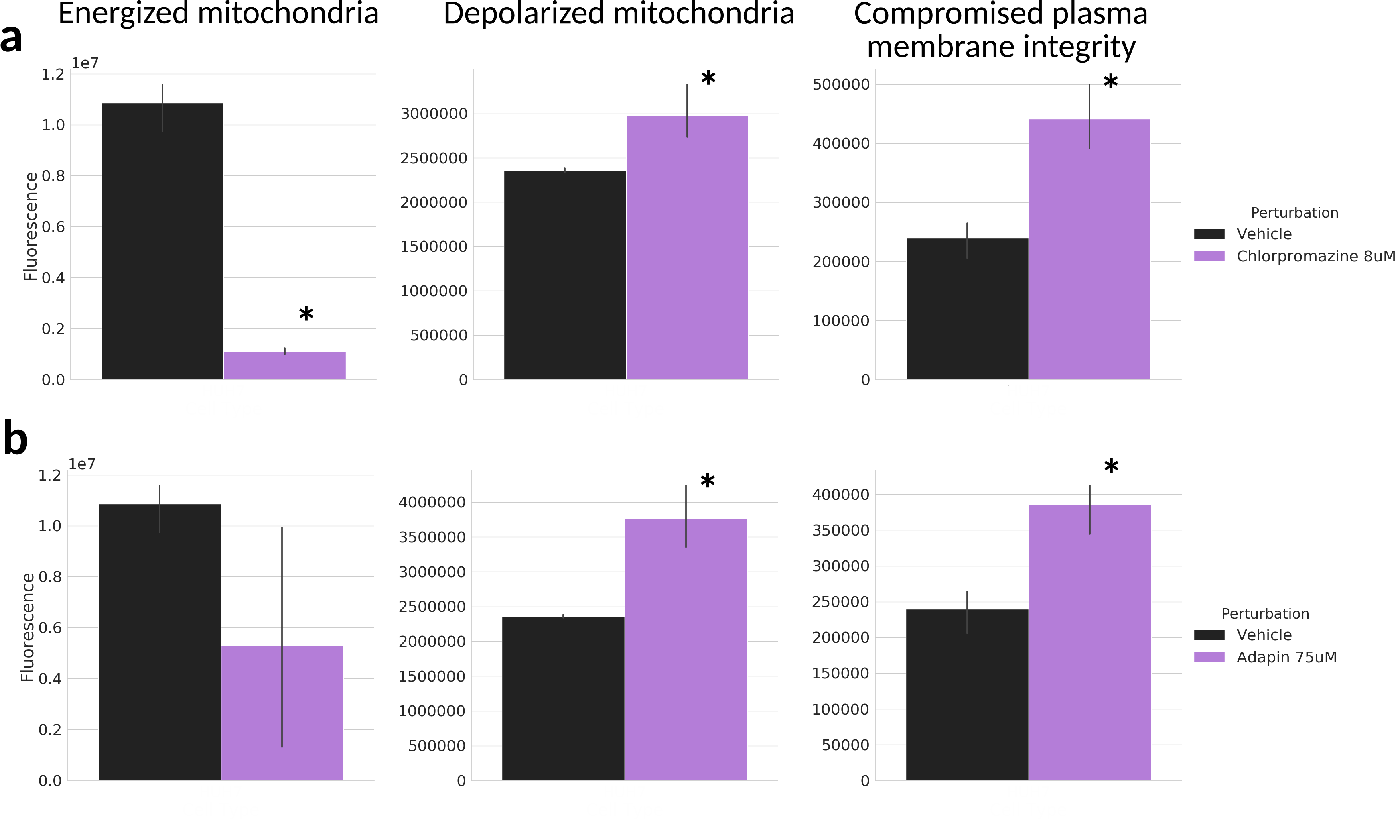
Experimental validation of MechSpy’s mechanistic prediction of mitochondrial toxicity for chlorpromazine and adapin. Fluorescense intensities of the three potential-sensitive MITO-ID dyes for HUH7 hepatocytes after 24 hour exposure to chlorpromazine (**a**) and adapin (**b**). The bars corresponding to treated cells marked with an asterisk (*) present a p-value smaller than 0.05 when compared to the untreated cells using a t-test.

The validation assays showed significant decrease in mitochondrial membrane potential and increased depolarization for HUH7 cells over prolonged exposure to 75*μ*M adapin (Fig. 7.a). This was also confirmed in a different hepatocyte cell line (HepG2, Supplemental Experimental Details). Adapin appeared to elicit a similar response in our control HCT116 cells, suggesting the mitochondrial-mediated toxicity may not be limited to hepatocytes. MechSpy’s prediction of mitochondrial toxicity was also verified for chlorpromazine, on HepG2 and particularly HUH7 cells where there was a significantly increased depolarization (Fig 7.b) when exposed to 8 a *μ*M concentration. In HCT116 cells there was a significant loss of mitochondrial membrane potential compared to vehicle control, yet not an increased membrane depolarization nor compromised membrane, suggesting chlorpromazine’s effects may be more tissue-specific.

We hope our mechanistic predictions for chemicals without an established mechanism of toxicity (summary in Table. 4, full list of predictions in Supplemental Table S3) can help guide further experimental work to seek hypothesis validation. Some interesting patterns were observed, such as hydroxyzine being enriched for the estrogen receptor-mediated mechanism (M8) in the two higher concentrations on hepatocytes. This is particularly interesting for hydroxyzine, as it has been linked to teratogenicity in rat models [49]. Many of these compounds exhibit common mechanistic predictions at different concentrations. Labetalol-exposed hepatocytes appear to primarily display oxidative stress (M4) as the most likely mechanism at toxic concentrations. In lung epithelial tissue, urea also appears to present a primarily oxidative stress (M4) mechanism. Nifedipine is a calcium channel antagonist that is actually administered to counter the toxicity of several other drugs, thus it’s unsurprising that in most cases MechSpy didn’t detect a significant enough enrichment of toxicity mechanisms. At the highest concentration exposed to hepatocytes, however, MechSpy predicted mitochondrial-mediated toxicity (M5) as the most likely mechanism, as well as the second most likely for treated kidney cells. The rare reported toxicity events associated to nifedipine are related to cardiotoxicity [50], which is consistent with mitochondria-mediated toxicity, since cardiomyocytes are highly energy-demanding cells and this adverse mechanism results in depletion of ATP.

**Table 4.**
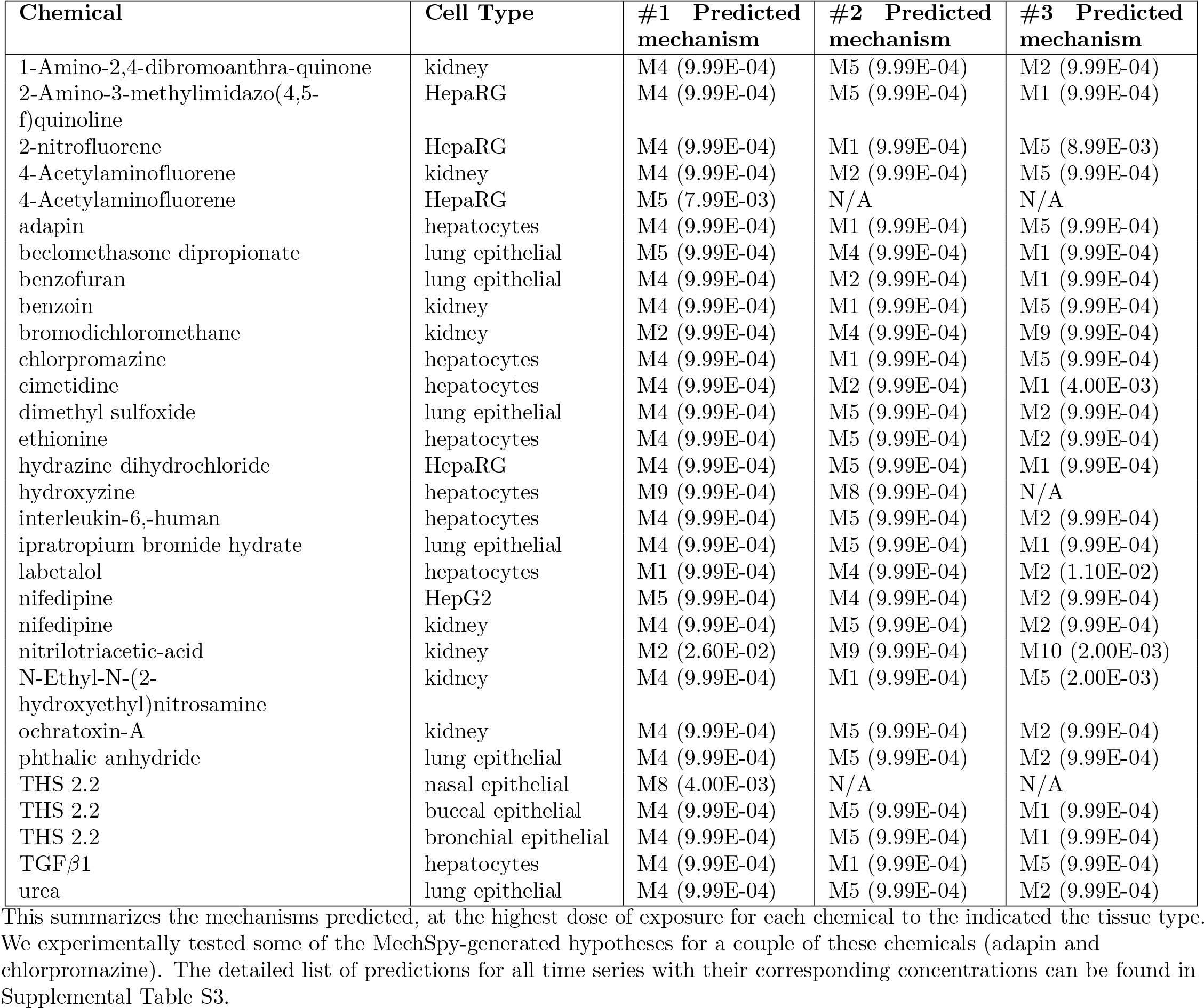
MechSpy predictions for chemicals for which there is no strong enough evidence of a particular mechanism of toxicity.

## Discussion

We present in this study a framework that holds great potential to aid the hypothesis generation process of mechanistic toxicology. Combining data from two sources, experimental results and existing knowledge, presents the best of both worlds: this is neither a purely data-driven inference without regards of context, nor a purely semantic knowledge-based exercise of what is plausible. Given the economic, practical and ethical burden in animal models to elucidate mechanisms of toxicity, MechSpy can also serve as a potential animal testing reduction or replacement tool. Its richness of knowledge sources makes it useful for data originating in other assays beyond gene expression, like proteomics, metabolomics, or chromatin accessibility. MechSpy has a direct application to both pre-clinical drug development and pharmacovigilance later on, to study rare side effects on subsets of the population. We demonstrate how using a coarse transcriptomics time series post-exposure to a known toxicant, our method can identify the most likely mechanisms among the top-three most strongly enriched.

The predictions generated by MechSpy are in agreement with the literature for many of the chemicals we tested at different doses (see Results), even when many of the expected mechanisms came from animal models across a variety of species, sometimes using different tissue types. We focused on the top-three most highly enriched mechanisms not only because this is a hypothesis generation aid, but also because in many cases there is no single mechanism of toxicity for a compound. Such is the case of acetaminophen (APAP), which has been known to elicit a toxic response via at least three different mechanisms [51]. Moreover, a compound’s mechanism of toxicity may depend on the dose of exposure. Some of the predicted mechanisms for those chemicals without an established mechanism of toxicity were validated experimentally, which shows a mechanistic inference framework like MechSpy is a robust and practical tool.

Some mechanisms from the curated list may be hard to detect from gene expression alone. Such is the case of M6, the cell cycle disruption one, which may require very narrow experimental time point margins to really detect it, or a different kind of assay altogether. The fact that this particular mechanism (M6) was missed for all compounds evaluated, may highlight the fact that either the long inter-sampling times or the assay types are inappropriate to identify it. As can be observed in Supplemental Table S2, several of the assays failing to predict any of the expected mechanisms among the top-three correspond to the lowest doses of exposure for that chemical. For these cases, the dose may not elicit any toxic response at all, or the changes in expression may be too subtle for MechSpy to detect them. Other chemicals may pose relatively low risk of toxicity to humans, like the case of coumarin, which proved hard to predict accurately and may require a much larger dose to elicit the expected oxidative stress response, extrapolated from animal models (a complex task, since coumarin presents different mechanisms of toxicity in mice, rats and humans [19]).

Not all exposure experiments to toxic compounds resulted in the expected predictions. Dibenz[a,h]anthracene is a particular case, given it’s generally known to result in an adverse response after exposure to ultraviolet (UV) light, leading to phototoxicity. Our evaluation was performed on exposure assays to lung epithelial cells without UV, which could potentially elucidate a different mechanism of toxicity. For methyltestosterone-treated cells, the choice of tissue (hepatocytes) could be a reason these assays were not enriched for estrogen receptor-mediated mechanisms.

A challenge we faced in this study was the availability of public datasets that used chemicals known to elicit every mechanism of toxicity we described. Despite the lack of public datasets involving chemicals known to act via mechanisms mediated by the aryl hydrocarbon receptor (M9) or androgen receptor (M10), we still included them in the process to show that these were not incorrectly predicted among most of the top-three results. All datasets consisted of microarray transcriptomics assays, which don’t necessarily have the best dynamic range of signal readouts, therefore other types of experiments like RNA-seq would be preferred. Furthermore, there may be certain mechanisms of toxicity that can only be detected using other assays than gene expression which, in the end, is a steady-state measurement of mature RNA, rather than a point-in-time measurement of transcription.

An inherent challenge of any kind of mechanistic study is also the dependence on the correct choices of time points and dosage. Multiple concentrations of the exposure dose must be tested, as cells could respond via different mechanisms. Such is the case of Cyclosporin A, which can elicit a toxic response via caspase-mediated apoptosis but, if the dose is high enough, then oxidative stress becomes the main reason for tissue damage [19]. Furthermore, several of the doses used in the datasets displayed in the results (Supplemental Table) are likely not high enough to elicit a toxic response. Unsurprisingly, the concentration of exposure was critical to start detecting strong enrichment of the expected mechanisms of some chemicals. Some illustrative examples were allopurinol, aspirin, coumarin, imipramine, or N-methyl-N-nitrosurea, for which MechSpy started predicting the expected mechanisms only at the highest concentration.

We took a conservative approach building the graph, using only human-curated relations (i.e. edges), therefore it will be worth exploring the improvements that can be achieved by incorporating computationally-inferred edges. We also acknowledge that not all mechanisms of toxicity are necessarily linear pathways, and may feature branches or cycles in their representation. A modification of the presented algorithms to seek enrichment in mechanisms of these other configurations is a topic for future development. A natural next step to this work will be to seek enrichment from other types of time series than from gene expression data, and to use instead (or in combination with) nascent transcription, proteomics and metabolomics assays at matching time points. We could also apply MechSpy to a proper semantic representation of existing AOPs from AOPwiki [30] or Effectopedia [52], rather than high-level molecular mechanisms of toxicity. Therefore, an integration between MechSpy and the AOPOntology [53] would also be desirable to explore.

## Conclusion

We envision that this mechanistic inference framework can be applied beyond the scope of toxicology, into any other discipline with a rich enough background knowledge represented with ontologies. The framework we present in this study can be extended to include other mechanisms as well, as long as they can be defined in terms of ontology concepts. The application of MechSpy goes beyond safety assessment and novel drug development, and can also be used to identify small molecules to be used as cancer therapeutics, based on their toxicity mechanism. Moreover, this mechanistic inference framework spans not just to other problems in molecular biology, but even to disciplines outside of the biomedical realm.

## Supporting information

**Supplemental Table S1. Literature sources used to determine each mechanism label.**

**Supplemental Table S2. MechSpy predictions for all time series with two or more time points with significant gene expression changes.**

**Supplemental Table S3. MechSpy predictions for chemicals for which there is no strong enough evidence of a particular mechanism of toxicity.**

**Supplemental Experimental Details. Additional details of the experimental validation of MechSpy-generated mechanistic hypotheses for adapin and chlorpromazine.**

## Software availability

All the code from MechSpy to process the samples and perform mechanistic inference is publicly available at https://github.com/ignaciot/MechSpy. The original version of the knowledge graph [20] used in this study can be found at https://github.com/callahantiff/PheKnowLator/wiki.

## Acknowledgements

We would like to thank Dr. Jared Brown, Dr. Melanie Joy, Dr. Kristina Brooks, and Dr. Manisha Patel at the toxicology department of University of Colorado, Denver, Anschutz medical campus, for the insightful discussions on mechanistic toxicology and feedback on the curated mechanisms. We are also very grateful for Dr. Jared Brown’s and Dr. Kristofer Fritz’s help with obtaining HepG2 cells for our validation assays.

## Author contributions

IJT: Mechanistic inference framework conceptualization and implementation, transcriptomics data processing, AOPwiki-derived edges in the KG, and curation of mechanisms. TC: Knowledge graph design and implementation, deductive closure. JTW: Experimental validation of our predictions for some compounds without established mechanisms of toxicity. NSM: Literature review to label mechanisms of toxicity for chemicals beyond Boelsterli’s textbook. RDD, LEH: Research supervision. IJT, TC, JTVW, RDD, and LEH contributed to the writing of this manuscript.

