## Supplementary material for "Applying knowledge-driven mechanistic inference to toxicogenomics": supplemental_experimental_details.pdf

### Experimental details for "Mechanistic inference from knowledge representation: a toxicogenomics case study"

Cells were treated with three concentrations and evaluated at two times points per concentration. Huh7s hepatocyte-derived carcinoma cell line, HepG2 hepatocyte-derived carcinoma cell line, and HCT116 human colorectal carcinoma cell line were grown and maintained in DMEM supplemented with 10% fetal bovine serum, 2 M L-glutamine and 1% Penicillin-Streptomycin and maintained at 37C, 5% $CO_2$. For the MMP assay, $2 \times 10^4$ cells were plated in triplicates in 96-well cell culture plates and incubated overnight. For chlorpromazine, cells were treated with 0.8 M, 4 M, and 8 M chlorpromazine hydrochloride (Sigma C8138) for 2 hr, 8 hr, and 24 hr. For adapin, cells were treated with 3 M, 15 M, and 75 M adapin (doxepin hydrochloride, Sigma D4526) or chloroform (as vehicle control) for 2 hr, 8 hr, and 24 hr. Mitochondrial membrane potential was measured using the Mito-ID membrane potential cytotoxicity kit (Enzo Life Sciences ENZ-51018-0025) following manufacturer protocol. The positive control, carbonyl cyanide 3-chlorophenylhydrazone (CCCP, 1-4$\mu$M) was added 15 minutes before the potential-sensitive dye was dispensed into each well. Sample wells were aspirated, washed with 1x Assay Solution, and 100 L Dual Detection Reagent Solution was added. The plates were incubated for 15 minutes at 37C, 5% $CO_2$. Dye fluorescence was quantified on a microplate reader (Molecular Device SpectraMax iD3) at three settings; ex 540 / em 570, ex 546 / em 674, ex 490 / em 590.

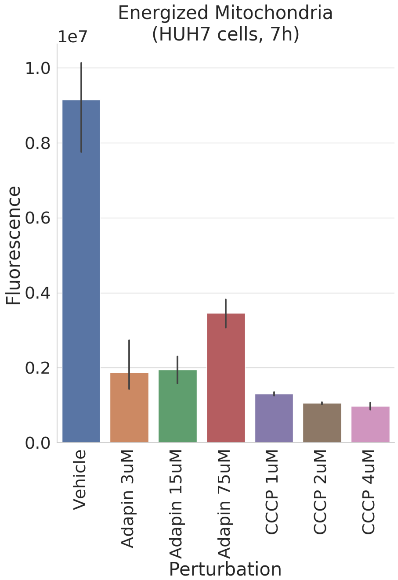


Figure 1: **Mitochondrial membrane potential for adapin on HUH7 cells after 7 hours of exposure.** Adapin at three concentrations is compared to CCCP at three concentrations, and untreated (vehicle) cells (n=3). Adapin-treated HUH7 cells show a reduction in energized mitochondria relative to negative control (Vehicle) that is on par with the provided positive control (CCCP).

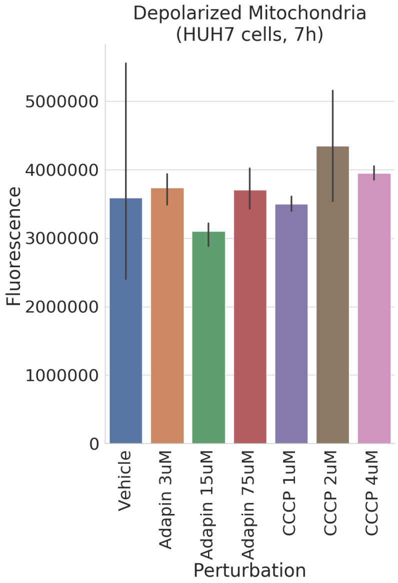


Figure 2: **Mitochondrial membrane depolarization for adapin on HUH7 cells after 7 hours of exposure.** Adapin-treated HUH7 cells don't show a significant increase in membrane depolarization at this time point relative to negative control (Vehicle), as neither does the positive control (CCCP). Axis labels as in Fig. 1 (n=3).

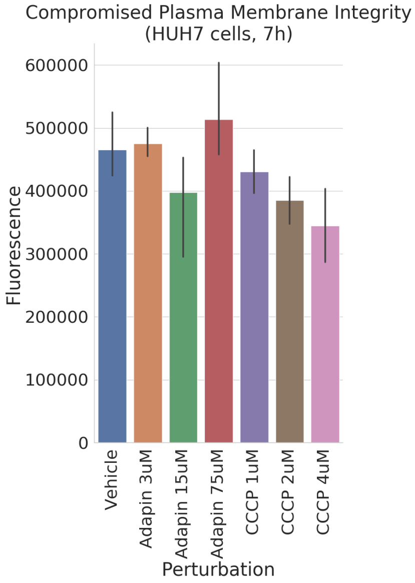


Figure 3: **Plasma membrane integrity for adapin on HUH7 cells after 7 hours of exposure.** Adapin-treated HUH7 cells don't show a significant increase in compromised membrane integrity relative to the negative control (Vehicle) or positive control (CCCP). Axis labels as in Fig. 1 (n=3).

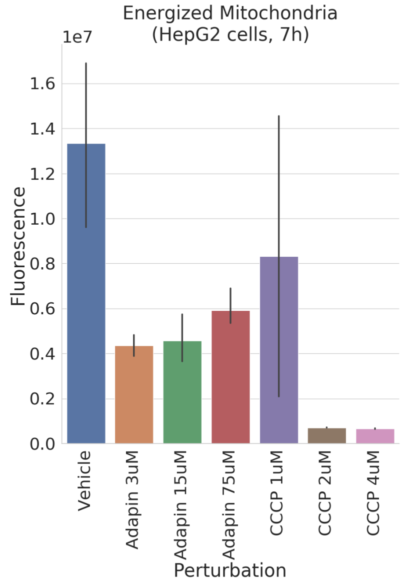


Figure 4: **Mitochondrial membrane potential for adapin on HepG2 cells after 7 hours of exposure.** Adapin-treated HepG2 cells show a reduction in energized mitochondria relative to negative control (Vehicle), following the same trend as the provided positive control (CCCP). Axis labels as in Fig. 1 (n=3).

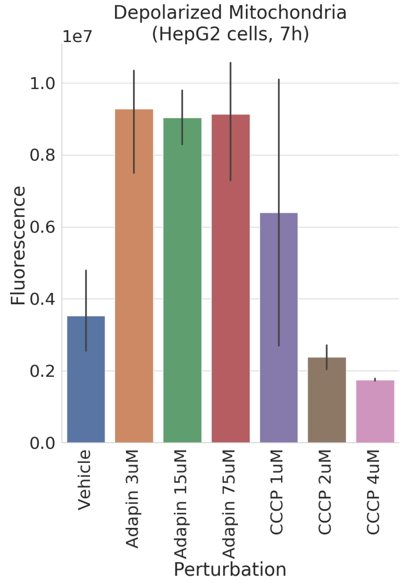


Figure 5: **Mitochondrial membrane depolarization for adapin on HepG2 cells after 7 hours of exposure.** Adapin-treated HepG2 cells show a significant increase in membrane depolarization relative to negative control (Vehicle), notably more than the positive control (CCCP). Axis labels as in Fig. 1 (n=3).

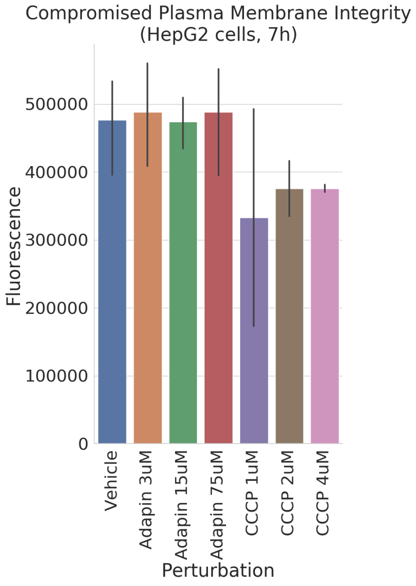


Figure 6: **Plasma membrane integrity for adapin on HepG2 cells after 7 hours of exposure.** No significant change in plasma membrane damage was observed in adapin-treated HepG2 cells relative to negative control (Vehicle). Axis labels as in Fig. 1 (n=3).

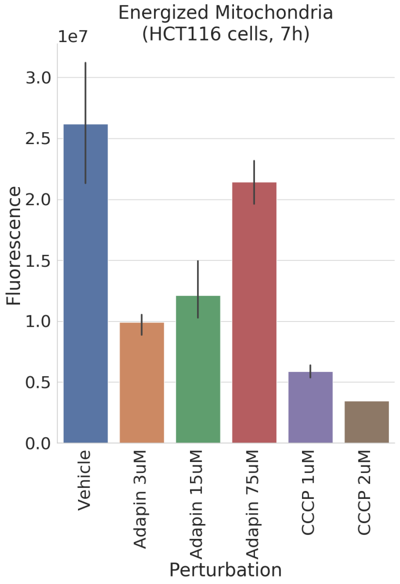


Figure 7: **Mitochondrial membrane potential for adapin on HCT116 cells after 7 hours of exposure.** Adapin at three concentrations is compared to CCCP at two concentrations, and untreated (vehicle) cells. Adapin-treated HCT116 cells show a reduction in energized mitochondria relative to negative control (Vehicle), following the same trend as the provided positive control (CCCP). (n=3).

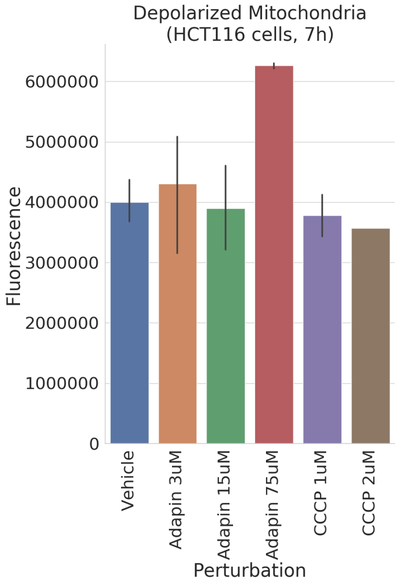


Figure 8: **Mitochondrial membrane depolarization for adapin on HCT116 cells after 7 hours of exposure.** Adapin-treated HCT116 cells show an increase in membrane depolarization at the highest dose tested (75$\mu$M) relative to negative control (Vehicle), and to positive control (CCCP). Axis labels as in Fig. 7 (n=3).

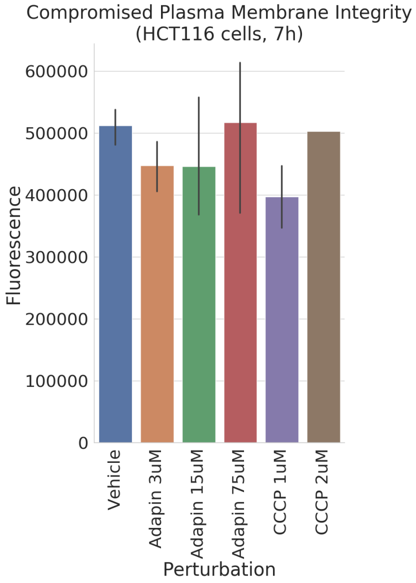


Figure 9: **Plasma membrane integrity for adapin on HCT116 cells after 7 hours of exposure.** No significant change in plasma membrane damage was observed in adapin-treated HCT116 cells at this time point, relative to negative control (Vehicle) or positive control (CCCP). Axis labels as in Fig. 7 (n=3).

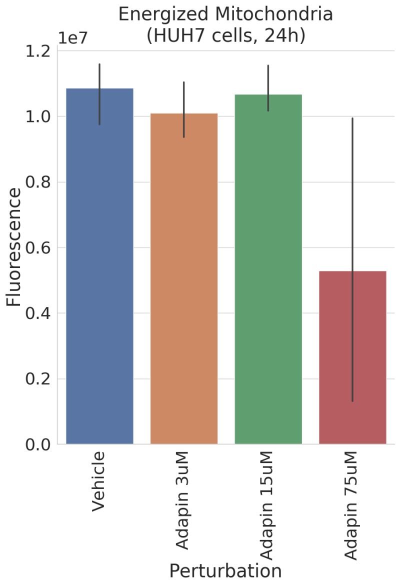


Figure 10: **Mitochondrial membrane potential for adapin on HUH7 cells after 24 hours of exposure.** Adapin at three concentrations is compared to untreated (vehicle) cells (n=3). Adapin-treated HUH7 cells at the highest dose tested (75$\mu$M) show a decrease in mitochondrial potential, relative to negative control (Vehicle). No difference to negative control is observed for the lower two doses.

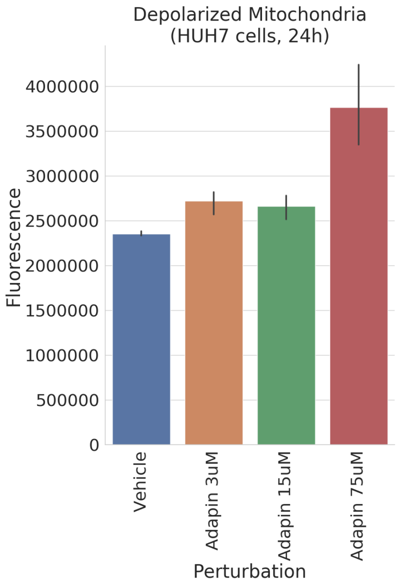


Figure 11: **Mitochondrial membrane depolarization for adapin on HUH7 cells after 24 hours of exposure.** Adapin-treated HUH7 cells at the highest dose tested (75$\mu$M) show an increase in membrane depolarization, relative to negative control (Vehicle). No difference to negative control is observed for the lower two doses. Axis labels as in Fig. 10 (n=3).

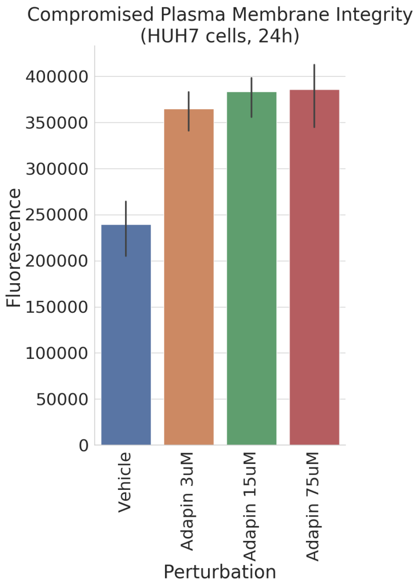


Figure 12: **Plasma membrane integrity for adapin on HUH7 cells after 24 hours of exposure.** Adapin-treated HCT116 cells show an increase in compromised plasma membrane integrity relative to negative control (Vehicle). Axis labels as in Fig. 10 (n=3).

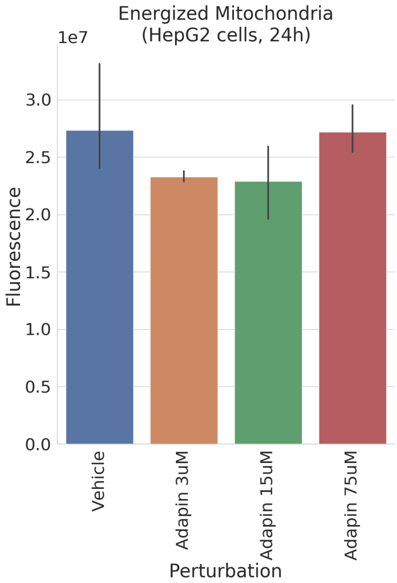


Figure 13: **Mitochondrial membrane potential for adapin on HepG2 cells after 24 hours of exposure.** Adapin-treated HepG2 cells show no notable difference in membrane potential relative to negative control (Vehicle). Axis labels as in Fig. 10 (n=3).

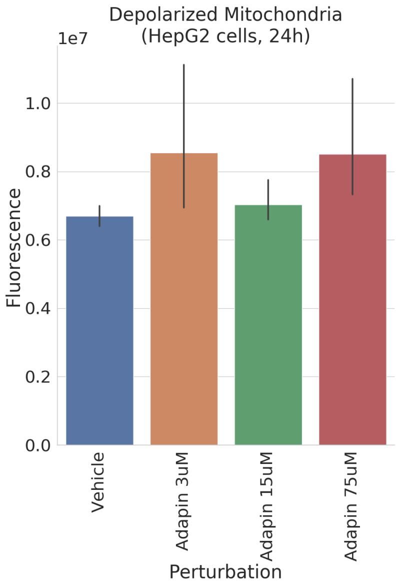


Figure 14: **Mitochondrial membrane depolarization for adapin on HepG2 cells after 24 hours of exposure.** Adapin-treated HepG2 cells begin to show a slight increase in membrane depolarization relative to negative control (Vehicle). Axis labels as in Fig. 10 (n=3).

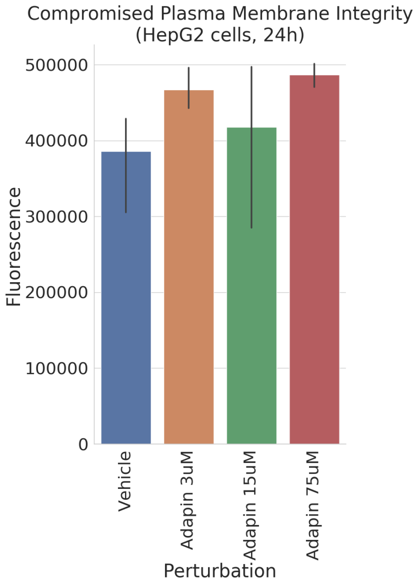


Figure 15: **Plasma membrane integrity for adapin on HepG2 cells after 24 hours of exposure.** Adapin-treated HepG2 cells show an increase in compromised plasma membrane integrity relative to negative control (Vehicle). Axis labels as in Fig. 10 (n=3).

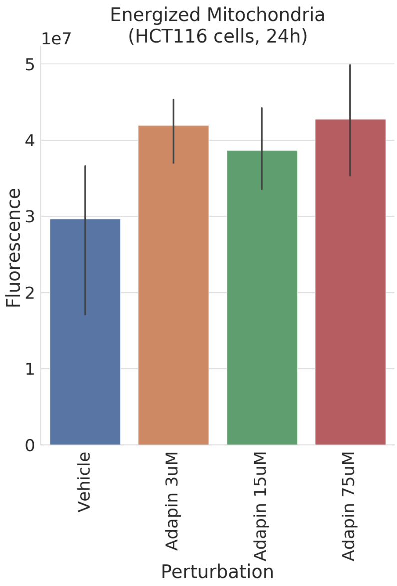


Figure 16: **Mitochondrial membrane potential for adapin on HCT116 cells after 24 hours of exposure.** Adapin-treated HCT116 cells surprisingly show an increase in membrane potential relative to negative control (Vehicle). Axis labels as in Fig. 10 (n=3).

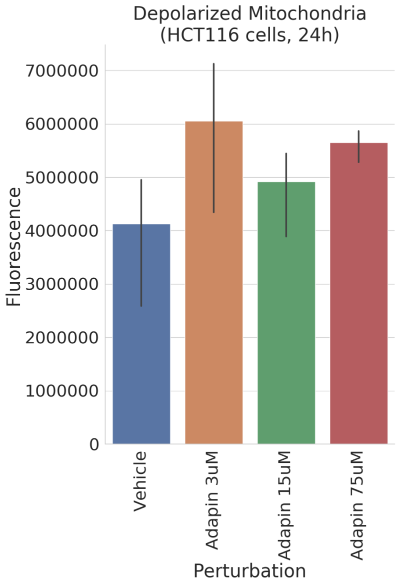


Figure 17: **Mitochondrial membrane depolarization for adapin on HCT116 cells after 24 hours of exposure.** Adapin-treated HCT116 cells show an increase in membrane depolarization relative to negative control (Vehicle). Axis labels as in Fig. 10 (n=3).

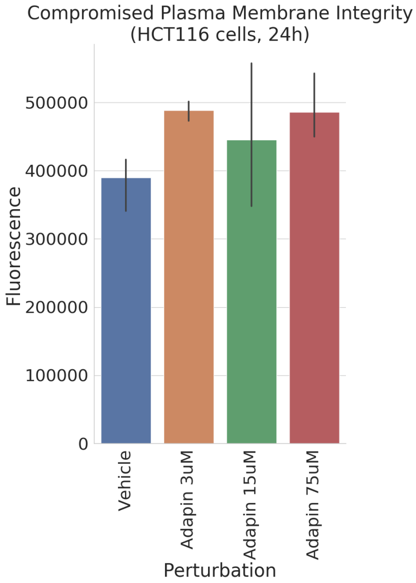


Figure 18: **Plasma membrane integrity for adapin on HCT116 cells after 24 hours of exposure.** Adapin-treated HCT116 cells show a slight increase in compromised plasma membrane integrity relative to negative control (Vehicle). Axis labels as in Fig. 10 (n=3).

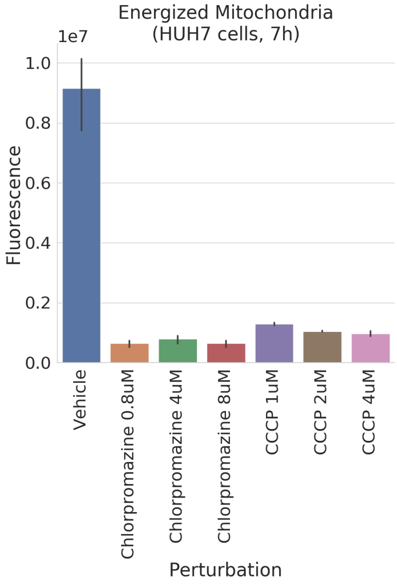


Figure 19: **Mitochondrial membrane potential for chlorpromazine on HUH7 cells after 7 hours of exposure.** Chlorpromazine at three concentrations is compared to CCCP at three concentrations, and untreated (vehicle) cells (n=2). Chlorpromazine-treated HUH7 cells show a decrease in membrane potential relative to negative control (Vehicle), on par with positive control (CCCP).

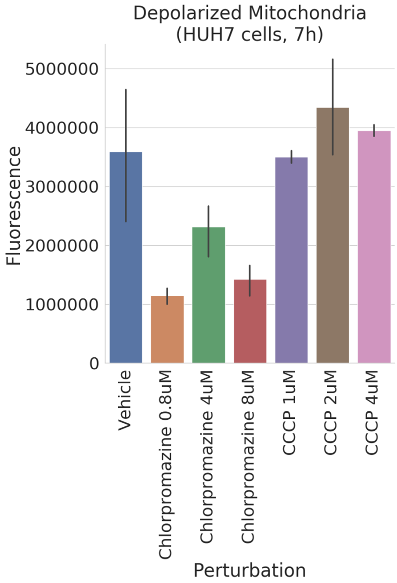


Figure 20: **Mitochondrial membrane depolarization for chlorpromazine on HUH7 cells after 7 hours of exposure.** Chlorpromazine-treated HUH7 cells show a decrease in membrane depolarization relative to negative control (Vehicle), unexpectedly lower than the positive control (CCCP). Axis labels as in Fig. 19 (n=2).

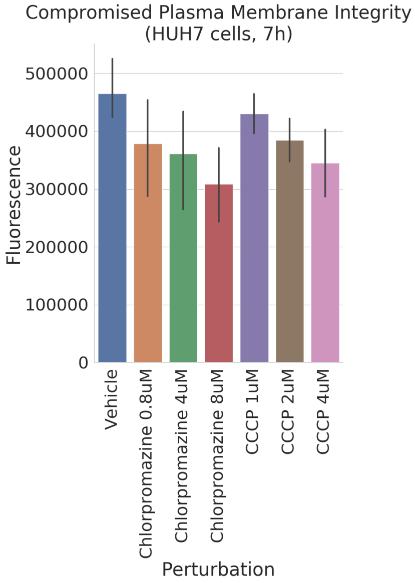


Figure 21: **Plasma membrane integrity for chlorpromazine on HUH7 cells after 7 hours of exposure.** Chlorpromazine-treated HUH7 cells show a slight decrease in compromised plasma membrane integrity relative to negative control (Vehicle), on par with positive control (CCCP). Axis labels as in Fig. 19 (n=2).

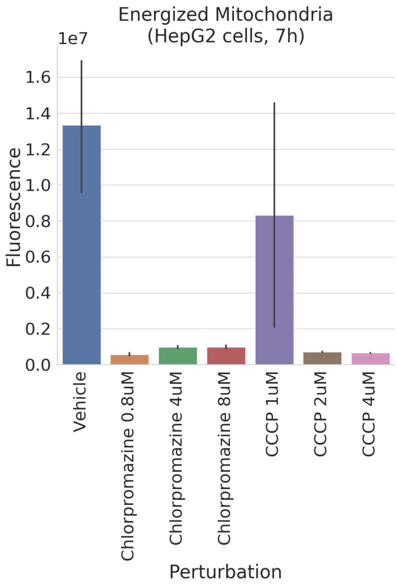


Figure 22: **Mitochondrial membrane potential for chlorpromazine on HepG2 cells after 7 hours of exposure.** Chlorpromazine-treated HepG2 cells show a decrease in membrane potential compared to negative control (Vehicle), on par with positive control (CCCP). Axis labels as in Fig. 19 (n=2).

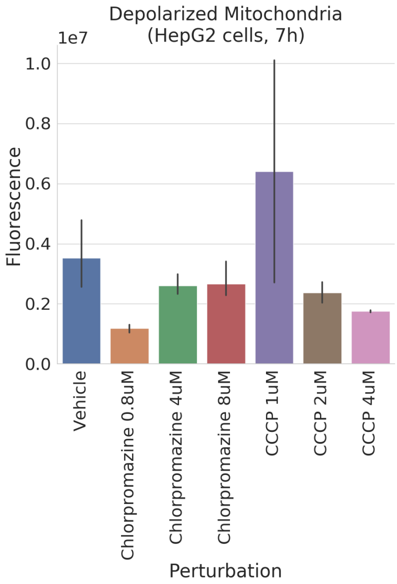


Figure 23: **Mitochondrial membrane depolarization for chlorpromazine on HepG2 cells after 7 hours of exposure.** Chlorpromazine-treated HepG2 cells don't show a notable increase of membrane depolarization at this time point relative to negative control (Vehicle), on par with the two highest doses of our positive control (CCCP). Axis labels as in Fig. 19 (n=2).

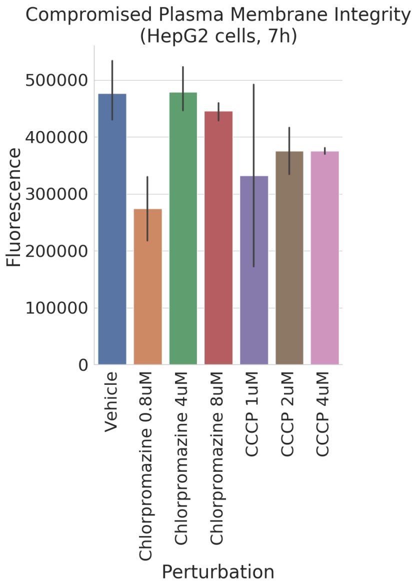


Figure 24: **Plasma membrane integrity for chlorpromazine on HepG2 cells after 7 hours of exposure.** No increase in compromised plasma membrane integrity is observed in chlorpromazine-treated HepG2 cells relative to negative control (Vehicle) at this time point. Axis labels as in Fig. 19 (n=2).

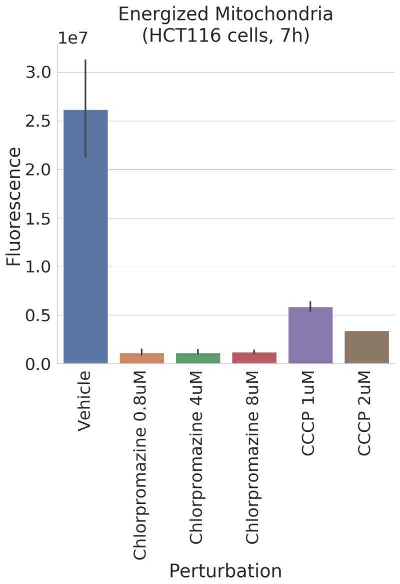


Figure 25: **Mitochondrial membrane potential for chlorpromazine on HCT116 cells after 7 hours of exposure.** Chlorpromazine at three concentrations is compared to CCCP at two concentrations, and untreated (vehicle) cells (n=2). Chlorpromazine-treated HCT116 cells shows a decrease in membrane potential relative to negative control (Vehicle), even lower than our positive control (CCCP).

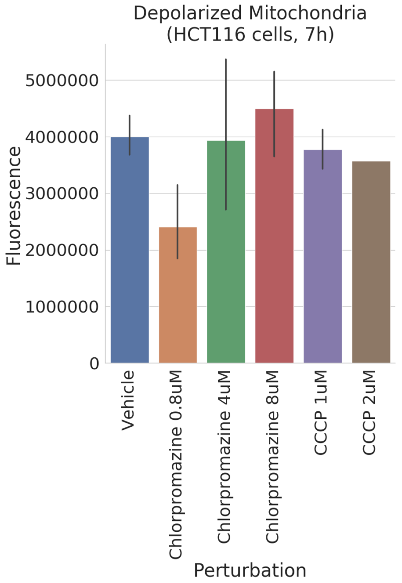


Figure 26: **Mitochondrial membrane depolarization for chlorpromazine on HCT116 cells after 7 hours of exposure.** Chlorpromazine-treated HCT116 cells show a slight increase of membrane depolarization at the highest dose tested (8$\mu$M), relative to negative control (Vehicle). Axis labels as in Fig. **??** (n=2).

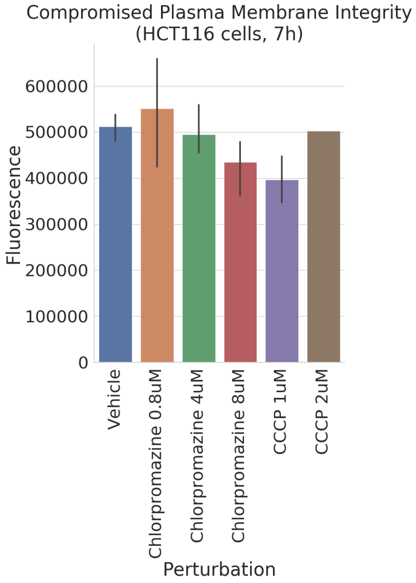


Figure 27: **Plasma membrane integrity for chlorpromazine on HCT116 cells after 7 hours of exposure.** No increase in compromised plasma membrane integrity is observed in chlorpromazine-treated HCT116 cells relative to negative control (Vehicle) at this time point. Axis labels as in Fig. **??** (n=2).

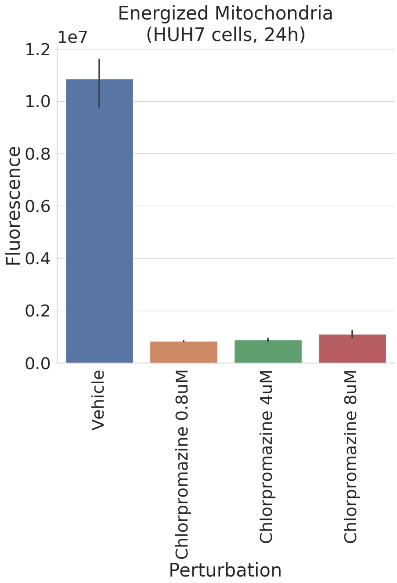


Figure 28: **Mitochondrial membrane potential for chlorpromazine on HUH7 cells after 24 hours of exposure.** Chlorpromazine at three concentrations is compared to untreated (vehicle) cells (n=3). Chlorpromazine-treated HUH7 cells show a decrease in membrane potential relative to negative control (Vehicle).

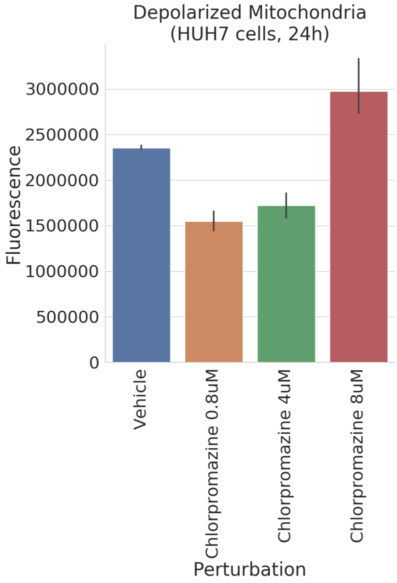


Figure 29: **Mitochondrial membrane depolarization for chlorpromazine on HUH7 cells after 24 hours of exposure.** Chlorpromazine-treated HUH7 cells show an increase in membrane depolarization at the highest dose tested (8$\mu$M), relative to negative control (Vehicle). Axis labels as in Fig. 28 (n=3).

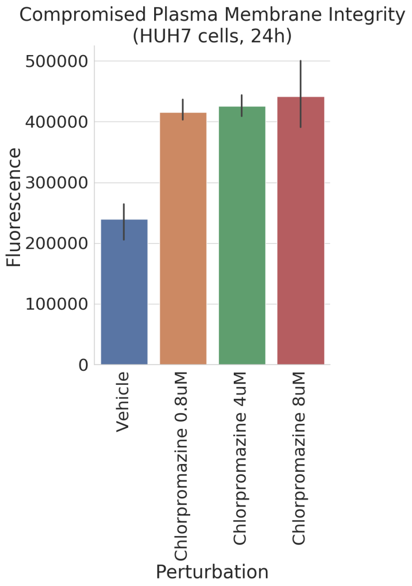


Figure 30: **Plasma membrane integrity for chlorpromazine on HUH7 cells after 24 hours of exposure.** Chlorpromazine-treated HUH7 cells show an increased compromised plasma membrane integrity relative to negative control (Vehicle). Axis labels as in Fig. 28 (n=3).

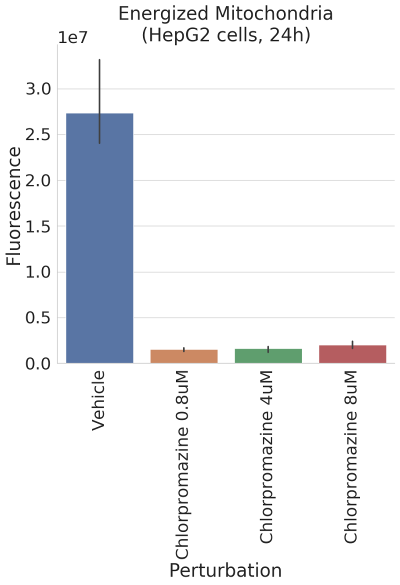


Figure 31: **Mitochondrial membrane potential for chlorpromazine on HepG2 cells after 24 hours of exposure.** Chlorpromazine-treated HepG2 cells show a decrease in membrane potential relative to negative control (Vehicle). Axis labels as in Fig. 28 (n=3).

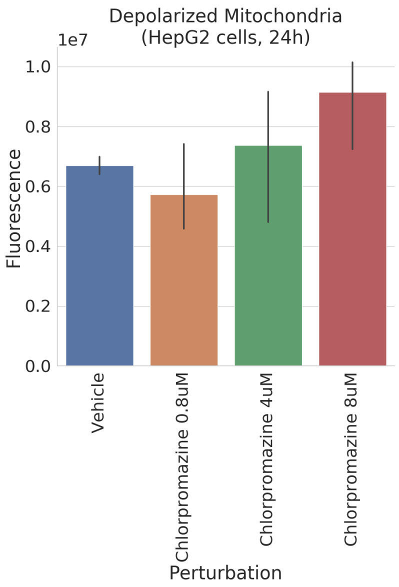


Figure 32: **Mitochondrial membrane depolarization for chlorpromazine on HepG2 cells after 24 hours of exposure.** Chlorpromazine-treated HepG2 cells show an increase in membrane depolarization at the highest dose tested (8$\mu$M), relative to negative control (Vehicle). Axis labels as in Fig. 28 (n=3).

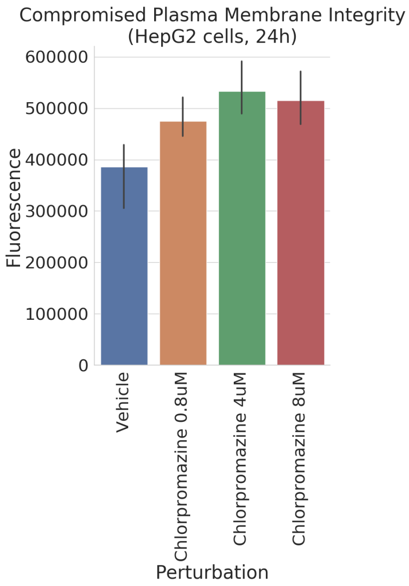


Figure 33: **Plasma membrane integrity for chlorpromazine on HepG2 cells after 24 hours of exposure.** Chlorpromazine-treated HepG2 cells show an increased compromised plasma membrane integrity relative to negative control (Vehicle). Axis labels as in Fig. 28 (n=3).

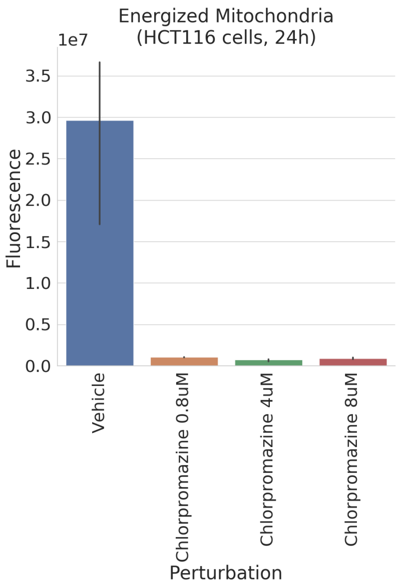


Figure 34: **Mitochondrial membrane potential for chlorpromazine on HCT116 cells after 24 hours of exposure.** Chlorpromazine-treated HCT116 cells show a decrease in membrane potential relative to negative control (Vehicle). Axis labels as in Fig. 28 (n=3).

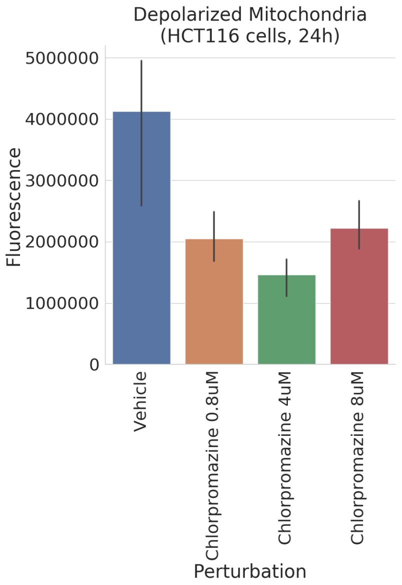


Figure 35: **Mitochondrial membrane depolarization for chlorpromazine on HCT116 cells after 24 hours of exposure.** Chlorpromazine-treated HepG2 cells show no increase in membrane depolarization relative to negative control (Vehicle), but rather the opposite. Axis labels as in Fig. 28 (n=3).

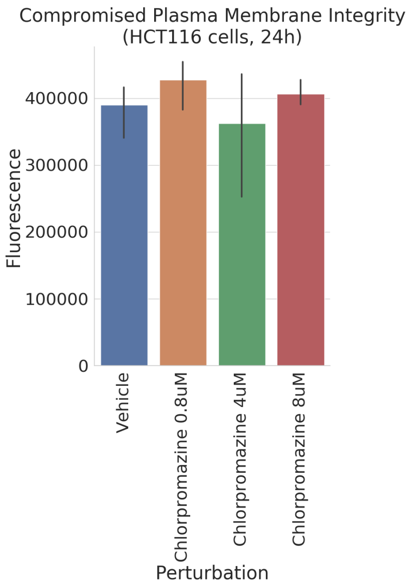


Figure 36: **Plasma membrane integrity for chlorpromazine on HCT116 cells after 24 hours of exposure.** No increase in compromised plasma membrane integrity is observed in chlorpromazine-treated HCT116 cells relative to negative control (Vehicle). Axis labels as in Fig. 28 (n=3).

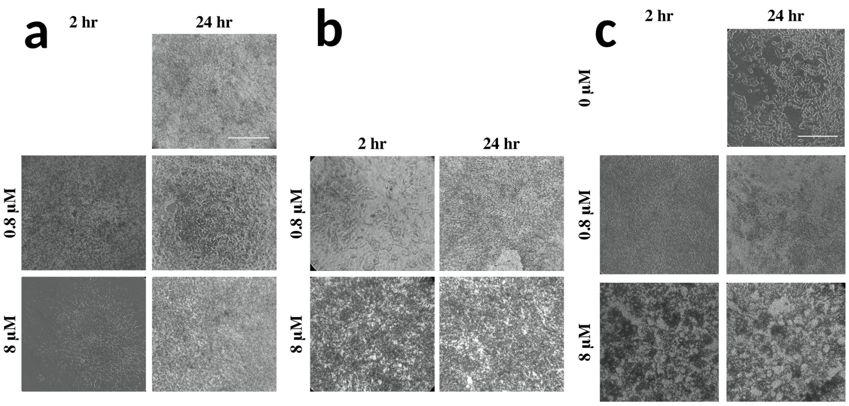


Figure 37: **Light microscopy of cells post-treatment with chlorpromazine at different time points.** HUH7 hepatocyte-derived carcinoma cells (**a**), HepG2 hepatocyte-derived carcinoma cells (**b**), and HCT116 human colorectal carcinoma cells (**c**) were imaged at 2 and 24 hours with 0.8$\mu$M and 8$\mu$M chlorpromazine. Magnification, 10x. Scale bar, 100 M

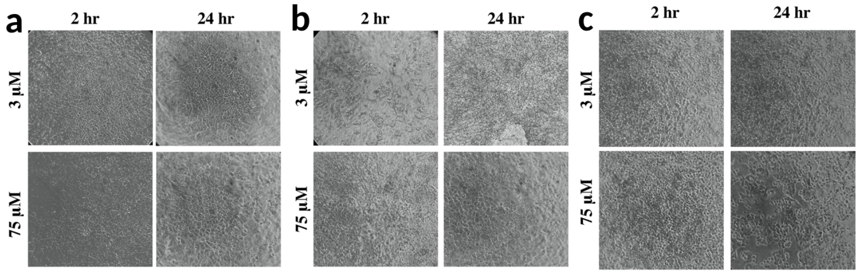


Figure 38: **Light microscopy of cells post-treatment with adapin at different time points.** HUH7 hepatocyte-derived carcinoma cells (**a**), HepG2 hepatocyte-derived carcinoma cells (**b**), and HCT116 human colorectal carcinoma cells (**c**) were imaged at 2 and 24 hours with 3$\mu$M and 75$\mu$M adapin. Magnification, 10x. Scale matching that of Fig. 37
