## Supplementary material for "Applying knowledge-driven mechanistic inference to toxicogenomics": supplemental_table_S1.pdf

**Literature sources of the various mechanism labels for each chemical.**

| Chemical | Mechanism Label | Source |
|---|---|---|
| 2-3-7-8-tetrachlorodibenzodioxine | M4 | Nie et al[1] |
| 2-3-7-8-tetrachlorodibenzodioxine | M9 | Boelsterli textbook[2] |
| 2-acetamidofluorene | M6 | Sparfel et al[3] |
| 4-(N-nitrosomethylamino)-1-(3-pyridyl)butan-1-one | M4 | Yalcin et al[4] |
| Diluted Mainstream Cigarette Smoke 3R4F | M4 | Lee et al[5], Poussin et al[6] |
| Diluted Mainstream Cigarette Smoke 3R4F | M7 | Lee et al[5] |
| Gallium arsenide | M4 | Flora et al[7] |
| N-Nitrosomorpholine | M5 | Korr et al[8] |
| N-methyl-N-nitrosurea | M3 | Furukawa et al[9] |
| N-methyl-N-nitrosurea | M4 | Furukawa et al[9] |
| S-(1,2-dichlorovinyl)-l-cysteine | M5 | Xu et al[10] |
| Tris-2-3-dibromopropyl-phosphate | M4 | Killilea et al[11] |
| acetaminophen | M2 | Boelsterli textbook[2] |
| acetaminophen | M4 | Boelsterli textbook[2] |
| acetaminophen | M5 | Hinson et al[12], McGill et al[13] |
| aflatoxin B1 | M4 | Boelsterli textbook[2] |
| allopurinol | M4 | Suzuki et al[14] |
| allyl alcohol | M2 | Brown et al[15] |
| allyl alcohol | M4 | Brown et al[15] |
| amiodarone | M5 | Felser et al[16] |
| aristolochic acid | M2 | Hsin et al[17] |
| aristolochic acid | M5 | Scorrano et al[18] |
| aspirin | M1 | Castaño et al[19], Hossain et al[20] |
| aspirin | M6 | Hossain et al[20], Ou et al[21] |
| azathioprine | M2 | Lee et al[22], Petit et al[23] |
| azathioprine | M5 | Lee et al[22], Petit et al[23] |
| benzbromarone | M5 | Felser et al[24] |
| benzo[a]pyrene | M4 | Boelsterli textbook[2] |
| benzyl alcohol | M1 | Chang et al[25] |
| bisphenol A | M5 | Bosch-Panadero et al[26] |
| bisphenol A | M8 | LaPensee et al[27] |
| bromobenzene | M5 | Wong et al[28] |
| cadmium dichloride | M4 | Skipper et al[29] |
| carbamazepine | M4 | Pirmohamed et al[30] |
| carbon tetrachloride | M4 | Boelsterli textbook[2] |
| citrinin | M1 | Salah et al[31] |
| citrinin | M7 | Salah et al[31] |
| clofibrate | M1 | Chen et al[32] |
| clofibrate | M7 | Chen et al[32] |
| clonidine | M1 | Fan et al[33] |

| | | |
|---|---|---|
| clonidine | M5 | Fan et al[33] |
| cobalt(2+) sulfate | M4 | Beyersmann et al[34] |
| coumarin | M4 | Boelsterli textbook[2] |
| cyclophosphamide hydrate | M1 | Iqubal et al[35] |
| cyclophosphamide hydrate | M2 | Iqubal et al[35] |
| cyclophosphamide hydrate | M4 | Omole et al[36] |
| cyclophosphamide hydrate | M5 | Iqubal et al[35] |
| cyclosporin A | M1 | Boelsterli textbook[2] |
| cyclosporin A | M5 | Wilmes et al[37] |
| diazepam | M6 | Chen et al[38] |
| dibenz[a,h]anthracene | M6 | Malik et al[39] |
| diclofenac sodium | M2 | Boelsterli textbook[2], Maiuri et al[40] |
| diclofenac sodium | M7 | Maiuri et al[40] |
| doxorubicin | M5 | Boelsterli textbook[2] |
| ethanol | M4 | Boelsterli textbook[2] |
| fluphenazine | M1 | Środa-Pomianek et al[41] |
| fluphenazine | M4 | Środa-Pomianek et al[41], Corte et al[42] |
| flutamide | M4 | Zhang et al[43] |
| flutamide | M5 | Zhang et al[43] |
| fumonisin B1 | M1 | Galvano et al[44] |
| fumonisin B1 | M4 | Singh et al[45] |
| fumonisin B1 | M7 | Singh et al[45] |
| gemfibrozil | M11 | Liu et al[46], Duez et al[47] |
| glibenclamide | M2 | Iwakura et al[48] |
| griseofulvin | M1 | Ho et al[49] |
| griseofulvin | M6 | Ho et al[49] |
| haloperidol | M1 | Hanagama et al[50] |
| haloperidol | M4 | Hanagama et al[50] |
| hexachlorobenzene | M1 | Śtarek-Wiechowicz et al[51] |
| hexachlorobenzene | M4 | Śtarek-Wiechowicz et al[51] |
| hydroquinone | M1 | Inayat-Hussain et al[52] |
| hydroquinone | M4 | Barreto et al[53] |
| ibuprofen | M5 | Al-Nasser[54] |
| ibuprofen | M6 | Elsisi et al[55] |
| imipramine | M1 | Piccotti et al[56] |
| imipramine | M11 | Piccotti et al[56] |
| indomethacin | M1 | Pantovic et al[57] |
| indomethacin | M4 | Pantovic et al[57] |
| interleukin-1-beta,-human | M1 | Takahashi et al[58] |
| interleukin-1-beta,-human | M1 | Takahashi et al[58] |
| isoniazid | M4 | Boelsterli textbook[2] |
| ketoconazole | M1 | Chen et al[59] |
| ketoconazole | M6 | Chen et al[59] |
| lead diacetate trihydrate | M4 | Fang et al[60] |
| lead diacetate trihydrate | M7 | Fang et al[60] |
| lomustine | M5 | Shinwari et al[61] |
| lomustine | M6 | Shinwari et al[61] |
| menthol | M5 | Berson et al[62] |
| methapyrilene hydrochloride | M1 | Mercer et al[63] |
| methapyrilene hydrochloride | M2 | Ratra et al[64] |
| methapyrilene hydrochloride | M5 | Ratra et al[64] |

| | | |
|---|---|---|
| methyltestosterone | M8 | Ma et al[65] |
| naphthyl isothiocyanate | M5 | Palmeira et al[66] |
| naphthyl-isothiocyanate | M4 | Palmeira et al[66] |
| nitrofurantoin | M4 | Tsuchiya et al[67] |
| nitrofurantoin | M5 | Omidi et al[68] |
| omeprazole | M4 | Seoane et al[69] |
| omeprazole | M5 | Seoane et al[69] |
| perhexiline | M5 | Oorts et al[70] |
| phenobarbital sodium | M4 | Boelsterli textbook[2] |
| phenylbutazone | M4 | Miura et al[71] |
| phenylbutazone | M5 | Yukiko et al[72] |
| phenytoin | M4 | Gallagher et al[73] |
| piperonyl butoxide | M4 | Muguruma et al[74] |
| pirinixic acid, WY-14643 | M2 | Taizo et al[75] |
| pirinixic acid, WY-14643 | M5 | Taizo et al[75] |
| potassium bromate | M4 | Ahmad et al[76] |
| propylthiouracil | M4 | Tang et al[77] |
| resorcinol | M5 | Skowroń et al[78] |
| rifampicin | M4 | Shen et al[79] |
| rifampicin | M11 | Huang et al[80] |
| rotenone | M5 | Boelsterli textbook[2] |
| sodium dichromate | M4 | Bagchi et al[81] |
| sodium metaarsenite | M4 | Oyagbemi et al[82] |
| streptozocin | M5 | Raza et al[83] |
| styrene | M4 | Röder-Stolinksi et al[84] |
| sulfasalazine | M4 | Linares et al[85] |
| tacrine | M5 | Boelsterli textbook[2] |
| tetracycline | M5 | Liu et al[86] |
| thioacetamide | M4 | Staňková et al[87] |
| thioacetamide | M5 | Amirtharaj et al[88] |
| thioridazine | M1 | Kang et al[89] |
| thioridazine | M6 | Kang et al[89] |
| thioridazine | M7 | Seervi et al[90] |
| tolbutamide | M2 | Smith et al[91], Efanova et al[92] |
| triclosan | M1 | Honkisz et al[93] |
| triclosan | M5 | Teplova et al[94] |
| valproic acid | M4 | Ramachandran et al[95] |
| valproic acid | M5 | Boelsterli textbook[2] |

Many of these mechanistic hypotheses come from a variety of animal models, examining a broad range of tissue types (often not the same tissue type than the publicly available experiments we used).

[68] Omidi M, Niknahad H, MohammadiBardbori A. Dithiothreitol (DTT) rescues mitochondria from nitrofurantoin-induced mitotoxicity in rat.

Journal of Biochemical and Molecular Toxicology. 2016;30(12):588–592. doi:10.1002/jbt.21825.

[80] Huang JH, Zhang C, Zhang DG, Li L, Chen X, Xu DX. Rifampicin-Induced Hepatic Lipid Accumulation: Association

with Up-Regulation of Peroxisome Proliferator-Activated Receptor in Mouse Liver. PLOS ONE. 2016;11(11):e0165787. doi:10.1371/journal.pone.0165787.
