## Supplementary material for "Applying knowledge-driven mechanistic inference to toxicogenomics": supplemental_table_S2.pdf

**Mechanistic inference results on all chemicals with compelling mechanistic explanations in the literature.**

| **Chemical** | **Dose** | **Cell Type** | **Known mechanisms** | **#1 Predicted mechanism** | **#2 Predicted mechanism** | **#3 Predicted mechanism** |
|---|---|---|---|---|---|---|
| 2,3,7,8-tetrachlorodibenzodioxine | 0.05$\mu$M | lung epithelial | M4, M6 | M5 (9.99E-04) | M1 (9.99E-04) | **M4** (7.99E-03) |
| 2,3,7,8-tetrachlorodibenzodioxine | 0.025$\mu$M | lung epithelial | M4, M6 | M5 (9.99E-04) | **M4** (9.99E-04) | M1 (9.99E-04) |
| 2-acetamidofluorene | 40.75$\mu$M | HepaRG | M6 | M5 (2.00E-03) | M8 (9.99E-04) | N/A |
| 3R4F (diluted mainstream cigarette smoke) | 0.2mgl | nasal epithelial | M4 | M5 (9.99E-04) | M1 (9.99E-04) | M2 (9.99E-04) |
| 3R4F (diluted mainstream cigarette smoke) | 0.15mgl | nasal epithelial | M4 | M5 (9.99E-04) | M8 (2.00E-03) | N/A |
| 3R4F (diluted mainstream cigarette smoke) | 0.25mgl | nasal epithelial | M4 | **M4** (9.99E-04) | M5 (9.99E-04) | M1 (9.99E-04) |
| 3R4F (diluted mainstream cigarette smoke) | 0.32mgl | buccal epithelial | M4 | **M4** (9.99E-04) | M5 (9.99E-04) | M1 (1.30E-02) |
| 3R4F (diluted mainstream cigarette smoke) | 0.51mgl | buccal epithelial | M4 | **M4** (9.99E-04) | M5 (9.99E-04) | M2 (9.99E-04) |
| 3R4F (total particulate matter from mainstream cigarette smoke) | 7.5$\mu$gl | bronchial epithelial | M4 | **M4** (9.99E-04) | M1 (9.99E-04) | M2 (9.99E-04) |

| | | | | | | |
|---|---|---|---|---|---|---|
| 4-(N-nitrosomethylamino)-1-(3-pyridyl)butan-1-one | 3750$\mu$M | HepaRG | M4 | **M4** (9.99E-04) | M5 (3.00E-03) | M2 (9.99E-04) |
| 4-(N-nitrosomethylamino)-1-(3-pyridyl)butan-1-one | 5000$\mu$M | HepG2 | M4 | **M4** (9.99E-04) | M2 (9.99E-04) | M1 (9.99E-04) |
| 4-(N-nitrosomethylamino)-1-(3-pyridyl)butan-1-one | 7500$\mu$M | HepaRG | M4 | **M4** (9.99E-04) | M5 (9.99E-04) | M1 (9.99E-04) |
| 4-(N-nitrosomethylamino)-1-(3-pyridyl)butan-1-one | 10000$\mu$M | HepG2 | M4 | M5 (9.99E-04) | **M4** (9.99E-04) | M1 (9.99E-04) |
| acetaminophen | 200$\mu$M | hepatocytes | M2, M4, M5 | M1 (6.99E-03) | **M4** (8.99E-03) | N/A |
| acetaminophen | 500$\mu$M | HepG2 | M2, M4, M5 | **M4** (9.99E-04) | **M2** (9.99E-04) | M1 (9.99E-04) |
| acetaminophen | 1000$\mu$M | hepatocytes | M2, M4, M5 | **M4** (9.99E-04) | **M5** (9.99E-04) | M1 (9.99E-04) |
| acetaminophen | 5000$\mu$M | hepatocytes | M2, M4, M5 | **M4** (9.99E-04) | **M2** (9.99E-04) | M1 (9.99E-04) |
| acetaminophen | 10000$\mu$M | HepG2 | M2, M4, M5 | **M4** (9.99E-04) | **M5** (9.99E-04) | **M2** (9.99E-04) |
| aflatoxin B1 | 0.24$\mu$M | hepatocytes | M4 | M5 (9.99E-04) | M1 (9.99E-04) | M2 (9.99E-04) |
| aflatoxin B1 | 1.2$\mu$M | hepatocytes | M4 | **M4** (9.99E-04) | M5 (9.99E-04) | M1 (9.99E-04) |
| aflatoxin B1 | 6$\mu$M | hepatocytes | M4 | **M4** (9.99E-04) | M5 (9.99E-04) | M1 (9.99E-04) |
| aflatoxin B1 | 1.6$\mu$M | HepG2 | M4 | **M4** (9.99E-04) | M5 (9.99E-04) | M1 (9.99E-04) |
| aflatoxin B1 | 2.5$\mu$M | HepaRG | M4 | **M4** (9.99E-04) | M1 (9.99E-04) | M5 (9.99E-04) |
| allopurinol | 5.6$\mu$M | hepatocytes | M4 | M5 (9.99E-04) | M2 (9.99E-04) | M8 (3.70E-02) |
| allopurinol | 28$\mu$M | hepatocytes | M4 | M5 (9.99E-04) | M2 (9.99E-04) | M1 (9.99E-04) |
| allopurinol | 140$\mu$M | hepatocytes | M4 | M5 (9.99E-04) | **M4** (9.99E-04) | M2 (9.99E-04) |
| allyl alcohol | 2.8$\mu$M | hepatocytes | M2, M4 | N/A | N/A | N/A |
| allyl alcohol | 14$\mu$M | hepatocytes | M2, M4 | M5 (9.99E-04) | **M2** (9.99E-04) | M1 (2.00E-02) |

| | | | | | | |
|---|---|---|---|---|---|---|
| allyl alcohol | 70$\mu$M | hepatocytes | M2, M4 | **M4** (9.99E-04) | M1 (9.99E-04) | M5 (9.99E-04) |
| amiodarone | 0.28$\mu$M | hepatocytes | M5 | N/A | N/A | N/A |
| amiodarone | 1.4$\mu$M | hepatocytes | M5 | N/A | N/A | N/A |
| amiodarone | 7$\mu$M | hepatocytes | M5 | **M5** (9.99E-04) | M4 (9.99E-04) | M2 (9.99E-04) |
| amiodarone | 7.5$\mu$M | hepatocytes | M5 | M4 (9.99E-04) | M1 (2.60E-02) | M7 (9.99E-04) |
| amiodarone | 15$\mu$M | hepatocytes | M5 | M4 (9.99E-04) | **M5** (9.99E-04) | M2 (5.99E-03) |
| aristolochic acid | 100$\mu$M | kidney | M1, M2, M5, M7 | M4 (9.99E-04) | **M5** (9.99E-04) | **M2** (9.99E-04) |
| aspirin | 600$\mu$M | hepatocytes | M1, M6 | M4 (9.99E-04) | M5 (3.00E-03) | M2 (9.99E-04) |
| aspirin | 3000$\mu$M | hepatocytes | M1, M6 | M4 (9.99E-04) | M2 (9.99E-04) | **M1** (9.99E-04) |
| azathioprine | 2.9$\mu$M | hepatocytes | M2, M5 | M4 (9.99E-04) | M1 (9.99E-04) | **M5** (9.99E-04) |
| azathioprine | 14.6$\mu$M | hepatocytes | M2, M5 | M1 (9.99E-04) | M4 (9.99E-04) | **M5** (9.99E-03) |
| azathioprine | 72.8$\mu$M | hepatocytes | M2, M5 | M1 (9.99E-04) | M4 (9.99E-04) | **M2** (9.99E-04) |
| benzbromarone | 4$\mu$M | hepatocytes | M5 | M4 (9.99E-04) | M2 (9.99E-04) | M1 (4.00E-03) |
| benzbromarone | 20$\mu$M | hepatocytes | M5 | M1 (9.99E-04) | M4 (9.99E-04) | **M5** (9.99E-04) |
| benzbromarone | 100$\mu$M | hepatocytes | M5 | M4 (9.99E-04) | M1 (9.99E-04) | **M5** (9.99E-04) |
| benzo[a]pyrene | 1$\mu$M | HepG2 | M4 | M5 (9.99E-04) | M9 (9.99E-04) | M8 (9.99E-04) |
| benzo[a]pyrene | 2$\mu$M | HepG2 | M4 | N/A | N/A | N/A |
| benzo[a]pyrene | 2.5$\mu$M | HepaRG | M4 | N/A | N/A | N/A |
| benzo[a]pyrene | 5$\mu$M | HepaRG | M4 | **M4** (9.99E-04) | M1 (9.99E-04) | M5 (7.99E-03) |
| benzo[a]pyrene | 10$\mu$M | lung epithelial | M4 | M5 (9.99E-04) | **M4** (9.99E-04) | M1 (9.99E-04) |
| benzo[a]pyrene | 20$\mu$M | lung epithelial | M4 | **M4** (9.99E-04) | M5 (9.99E-04) | M2 (9.99E-04) |
| benzo[a]pyrene | 50$\mu$M | kidney | M4 | **M4** (9.99E-04) | M5 (9.99E-04) | M2 (9.99E-04) |
| benzyl alcohol | 10mM | kidney | M1 | **M1** (9.99E-04) | M4 (9.99E-04) | M5 (9.99E-04) |
| bisphenol A | 12.5$\mu$M | lung epithelial | M5 | M4 (9.99E-04) | **M5** (9.99E-04) | M2 (9.99E-04) |
| bisphenol A | 25$\mu$M | lung epithelial | M5 | M4 (9.99E-04) | **M5** (9.99E-04) | M2 (9.99E-04) |
| bromobenzene | 40$\mu$M | hepatocytes | M5 | **M5** (2.90E-02) | N/A | N/A |

| | | | | | | |
|---|---|---|---|---|---|---|
| bromobenzene | 200$\mu$M | hepatocytes | M5 | M1 (5.00E-03) | M4 (2.00E-03) | M3 (9.99E-03) |
| cadmium dichloride | 6.25$\mu$M | lung epithelial | M4 | M5 (9.99E-04) | **M4** (9.99E-04) | M2 (9.99E-04) |
| cadmium dichloride | 12.5$\mu$M | lung epithelial | M4 | **M4** (9.99E-04) | M1 (9.99E-04) | M2 (9.99E-04) |
| carbamazepine | 12$\mu$M | hepatocytes | M4 | M5 (2.60E-02) | M2 (9.99E-04) | M7 (1.20E-02) |
| carbamazepine | 60$\mu$M | hepatocytes | M4 | **M4** (9.99E-04) | M2 (9.99E-04) | M1 (9.99E-04) |
| carbamazepine | 300$\mu$M | hepatocytes | M4 | **M4** (9.99E-04) | M5 (9.99E-04) | M2 (9.99E-04) |
| carbon tetrachloride | 300$\mu$M | hepatocytes | M4 | **M4** (9.99E-04) | M2 (9.99E-04) | M1 (9.99E-04) |
| carbon tetrachloride | 1500$\mu$M | hepatocytes | M4 | M5 (7.99E-03) | M1 (9.99E-04) | M9 (9.99E-04) |
| carbon tetrachloride | 7500$\mu$M | hepatocytes | M4 | M1 (3.00E-03) | **M4** (9.99E-04) | N/A |
| chlorothalonil | 100nM | kidney | M5 | M4 (9.99E-04) | M2 (9.99E-04) | M1 (9.99E-04) |
| citrinin | 66$\mu$M | kidney | M1, M7 | **M1** (1.60E-02) | N/A | N/A |
| clofibrate | 12$\mu$M | hepatocytes | M1, M7 | M5 (9.99E-04) | **M1** (5.00E-03) | M2 (9.99E-04) |
| clofibrate | 60$\mu$M | hepatocytes | M1, M7 | M4 (9.99E-04) | M5 (9.99E-04) | M2 (9.99E-04) |
| clofibrate | 300$\mu$M | hepatocytes | M1, M7 | M4 (9.99E-04) | **M1** (9.99E-04) | M2 (9.99E-04) |
| clonidine | 0.1$\mu$M | HepaRG | M1, M5 | M4 (9.99E-04) | **M5** (9.99E-04) | **M1** (9.99E-04) |
| clonidine | 0.0948$\mu$M | HepG2 | M1, M5 | M4 (9.99E-04) | **M5** (9.99E-04) | M2 (9.99E-04) |
| clonidine | 1000$\mu$M | kidney | M1, M5 | **M5** (9.99E-04) | M4 (9.99E-04) | **M1** (9.99E-04) |
| clonidine | 2500$\mu$M | kidney | M1, M5 | M4 (9.99E-04) | **M5** (9.99E-04) | M2 (9.99E-04) |
| cobalt(2+) sulfate | 75$\mu$M | lung epithelial | M4 | M5 (9.99E-04) | **M4** (9.99E-04) | M1 (9.99E-04) |
| cobalt(2+) sulfate | 150$\mu$M | lung epithelial | M4 | M5 (9.99E-04) | **M4** (9.99E-04) | M2 (9.99E-04) |
| coumarin | 12$\mu$M | hepatocytes | M4 | M5 (9.99E-04) | M3 (1.40E-02) | N/A |
| coumarin | 60$\mu$M | hepatocytes | M4 | M3 (9.99E-04) | N/A | N/A |
| coumarin | 300$\mu$M | hepatocytes | M4 | **M4** (9.99E-04) | M1 (9.99E-04) | M2 (9.99E-04) |
| cyclophosphamide | 80$\mu$M | hepatocytes | M1, M2, M4, M5 | **M5** (9.99E-04) | **M4** (9.99E-04) | **M1** (2.00E-03) |
| cyclophosphamide | 400$\mu$M | hepatocytes | M1, M2, M4, M5 | **M4** (9.99E-04) | **M1** (9.99E-04) | **M5** (9.99E-04) |

| | | | | | | |
|---|---|---|---|---|---|---|
| cyclophosphamide | 2000$\mu$M | hepatocytes | M1, M2, M4, M5 | **M4** (9.99E-04) | **M1** (9.99E-04) | **M5** (9.99E-04) |
| cyclophosphamide hydrate | 4.7$\mu$M | HepG2 | M1, M2, M4, M5 | **M5** (9.99E-04) | **M4** (9.99E-04) | **M1** (9.99E-04) |
| cyclophosphamide hydrate | 700$\mu$M | HepaRG | M1, M2, M4, M5 | **M4** (9.99E-04) | **M5** (9.99E-04) | **M1** (9.99E-04) |
| cyclophosphamide hydrate | 5000$\mu$M | lung epithelial | M1, M2, M4, M5 | **M5** (9.99E-04) | **M4** (9.99E-04) | **M2** (9.99E-04) |
| cyclophosphamide hydrate | 10000$\mu$M | lung epithelial | M1, M2, M4, M5 | **M4** (9.99E-04) | **M2** (9.99E-04) | **M1** (9.99E-04) |
| cyclosporin A | 1.2$\mu$M | hepatocytes | M1, M5 | **M5** (9.99E-04) | M4 (6.99E-03) | **M1** (2.00E-03) |
| cyclosporin A | 6$\mu$M | hepatocytes | M1, M5 | **M5** (9.99E-04) | **M1** (9.99E-04) | M2 (6.99E-03) |
| cyclosporin A | 3$\mu$M | hepatocytes | M1, M5 | M4 (9.99E-04) | **M5** (9.99E-04) | M2 (9.99E-04) |
| cyclosporin A | 20$\mu$M | hepatocytes | M1, M5 | M4 (9.99E-04) | **M5** (9.99E-04) | **M1** (9.99E-04) |
| diazepam | 10$\mu$M | hepatocytes | M6 | M1 (9.99E-04) | M4 (9.99E-04) | M5 (3.00E-03) |
| diazepam | 50$\mu$M | hepatocytes | M6 | M4 (9.99E-04) | M1 (9.99E-04) | M5 (9.99E-04) |
| diazepam | 250$\mu$M | hepatocytes | M6 | M4 (9.99E-04) | M5 (9.99E-04) | M2 (9.99E-04) |
| dibenz[a,h]anthracene | 62.5$\mu$M | lung epithelial | M6 | M4 (9.99E-04) | M5 (9.99E-04) | M2 (9.99E-04) |
| dibenz[a,h]anthracene | 125$\mu$M | lung epithelial | M6 | M4 (9.99E-04) | M5 (9.99E-04) | M2 (9.99E-04) |
| diclofenac | 16$\mu$M | hepatocytes | M2, M7 | **M2** (9.99E-04) | M4 (6.99E-03) | M8 (7.99E-03) |
| diclofenac | 80$\mu$M | hepatocytes | M2, M7 | M4 (9.99E-04) | M1 (9.99E-04) | M5 (9.99E-04) |
| diclofenac | 400$\mu$M | hepatocytes | M2, M7 | M4 (9.99E-04) | **M2** (9.99E-04) | M1 (9.99E-04) |
| diclofenac | 30$\mu$M | kidney | M2, M7 | M4 (9.99E-04) | M5 (9.99E-04) | **M2** (9.99E-04) |
| diclofenac | 40$\mu$M | HepG2 | M2, M7 | N/A | N/A | N/A |
| diclofenac | 80$\mu$M | HepG2 | M2, M7 | M4 (9.99E-04) | M1 (9.99E-04) | **M2** (9.99E-04) |
| doxorubicin | 0.4$\mu$M | hepatocytes | M5 | M1 (9.99E-04) | **M5** (6.99E-03) | M2 (9.99E-04) |
| doxorubicin | 2$\mu$M | hepatocytes | M5 | M4 (9.99E-04) | M1 (9.99E-04) | M2 (9.99E-04) |

| | | | | | | |
|---|---|---|---|---|---|---|
| doxorubicin | 10$\mu$M | hepatocytes | M5 | **M5** (9.99E-04) | M4 (9.99E-04) | M1 (9.99E-04) |
| ethanol | 10000$\mu$M | hepatocytes | M4 | M2 (9.99E-04) | **M4** (9.99E-04) | M1 (9.99E-04) |
| fluphenazine | 0.8$\mu$M | hepatocytes | M1, M4 | M5 (9.99E-04) | M2 (3.60E-02) | M8 (3.00E-03) |
| fluphenazine | 4$\mu$M | hepatocytes | M1, M4 | **M4** (9.99E-04) | M5 (9.99E-04) | M2 (9.99E-04) |
| fluphenazine | 20$\mu$M | hepatocytes | M1, M4 | **M4** (9.99E-04) | **M1** (9.99E-04) | M5 (2.00E-03) |
| flutamide | 2$\mu$M | hepatocytes | M4, M5 | **M5** (9.99E-04) | **M4** (9.99E-04) | M1 (4.60E-02) |
| flutamide | 10$\mu$M | hepatocytes | M4, M5 | **M4** (9.99E-04) | M2 (9.99E-04) | **M5** (9.99E-04) |
| flutamide | 50$\mu$M | hepatocytes | M4, M5 | **M4** (9.99E-04) | M1 (9.99E-04) | M2 (9.99E-04) |
| fumonisin B1 | 2.42mM | HepaRG | M1, M4, M7 | N/A | N/A | N/A |
| fumonisin B1 | 100$\mu$M | kidney | M1, M4, M7 | **M4** (9.99E-04) | M2 (9.99E-04) | M5 (2.50E-02) |
| gallium arsenide | 3.18$\mu$M | kidney | M4 | M1 (9.99E-04) | **M4** (9.99E-04) | M5 (9.99E-04) |
| gemfibrozil | 4$\mu$M | hepatocytes | M11 | M5 (9.99E-04) | M4 (9.99E-04) | M2 (9.99E-04) |
| gemfibrozil | 20$\mu$M | hepatocytes | M11 | M4 (9.99E-04) | M1 (5.00E-02) | M8 (3.00E-03) |
| gemfibrozil | 100$\mu$M | hepatocytes | M11 | M4 (9.99E-04) | M2 (9.99E-04) | M5 (9.99E-04) |
| glibenclamide | 0.8$\mu$M | hepatocytes | M2 | N/A | N/A | N/A |
| glibenclamide | 4$\mu$M | hepatocytes | M2 | M4 (9.99E-04) | M5 (3.00E-03) | M1 (3.00E-02) |
| glibenclamide | 20$\mu$M | hepatocytes | M2 | M4 (9.99E-04) | **M2** (9.99E-04) | M5 (2.00E-03) |
| griseofulvin | 0.8$\mu$M | hepatocytes | M1, M6 | **M1** (9.99E-04) | M2 (9.99E-04) | M5 (3.10E-02) |
| griseofulvin | 4$\mu$M | hepatocytes | M1, M6 | **M1** (2.80E-02) | M8 (9.99E-04) | N/A |
| griseofulvin | 20$\mu$M | hepatocytes | M1, M6 | M4 (9.99E-04) | M5 (9.99E-04) | M2 (9.99E-04) |
| haloperidol | 0.8$\mu$M | hepatocytes | M1, M4 | **M1** (9.99E-04) | **M4** (2.00E-03) | M3 (9.99E-04) |
| haloperidol | 4$\mu$M | hepatocytes | M1, M4 | **M1** (3.00E-03) | **M4** (2.40E-02) | M2 (9.99E-04) |
| haloperidol | 20$\mu$M | hepatocytes | M1, M4 | **M4** (9.99E-04) | M5 (9.99E-04) | M2 (9.99E-04) |
| hexachlorobenzene | 1.2$\mu$M | hepatocytes | M1, M4 | M5 (4.00E-02) | M8 (6.99E-03) | N/A |
| hexachlorobenzene | 6$\mu$M | hepatocytes | M1, M4 | M5 (9.99E-04) | M2 (9.99E-04) | M8 (9.99E-04) |
| hexachlorobenzene | 30$\mu$M | hepatocytes | M1, M4 | **M4** (9.99E-04) | M5 (9.99E-04) | M2 (9.99E-04) |
| hydroquinone | 252$\mu$M | kidney | M1, M4 | **M4** (9.99E-04) | M2 (9.99E-04) | M5 (9.99E-04) |

| | | | | | | |
|---|---|---|---|---|---|---|
| ibuprofen | 40$\mu$M | hepatocytes | M5, M6 | M1 (5.00E-03) | M2 (3.50E-02) | M9 (3.70E-02) |
| ibuprofen | 200$\mu$M | hepatocytes | M5, M6 | N/A | N/A | N/A |
| ibuprofen | 1000$\mu$M | hepatocytes | M5, M6 | M4 (4.00E-03) | M1 (2.90E-02) | M2 (1.80E-02) |
| imipramine | 4$\mu$M | hepatocytes | M1, M11 | M2 (4.00E-03) | N/A | N/A |
| imipramine | 20$\mu$M | hepatocytes | M1, M11 | N/A | N/A | N/A |
| imipramine | 100$\mu$M | hepatocytes | M1, M11 | M9 (7.99E-03) | M10 (2.00E-03) | **M11** (2.00E-02) |
| indomethacin | 8$\mu$M | hepatocytes | M1, M4 | N/A | N/A | N/A |
| indomethacin | 40$\mu$M | hepatocytes | M1, M4 | **M4** (9.99E-04) | **M1** (9.99E-04) | M5 (9.99E-04) |
| indomethacin | 200$\mu$M | hepatocytes | M1, M4 | **M4** (9.99E-04) | **M1** (9.99E-04) | M5 (4.70E-02) |
| interleukin 1 beta,-human | 2$\mu$M | hepatocytes | M1 | M4 (9.99E-04) | **M1** (9.99E-04) | M5 (2.00E-03) |
| interleukin 1 beta,-human | 10$\mu$M | hepatocytes | M1 | M4 (9.99E-04) | **M1** (9.99E-04) | M5 (9.99E-04) |
| interleukin 1 beta,-human | 50$\mu$M | hepatocytes | M1 | M4 (9.99E-04) | **M1** (9.99E-04) | M5 (2.00E-03) |
| isoniazid | 400$\mu$M | hepatocytes | M4 | **M4** (9.99E-04) | M2 (9.99E-04) | M1 (9.99E-04) |
| isoniazid | 2000$\mu$M | hepatocytes | M4 | **M4** (9.99E-04) | M2 (9.99E-04) | M5 (9.99E-04) |
| isoniazid | 10000$\mu$M | hepatocytes | M4 | **M4** (9.99E-04) | M2 (9.99E-04) | M5 (9.99E-04) |
| ketoconazole | 0.6$\mu$M | hepatocytes | M1, M6 | M5 (9.99E-04) | M2 (9.99E-04) | M8 (9.99E-04) |
| ketoconazole | 3$\mu$M | hepatocytes | M1, M6 | M4 (9.99E-04) | **M1** (9.99E-04) | M5 (9.99E-04) |
| ketoconazole | 15$\mu$M | hepatocytes | M1, M6 | **M1** (9.99E-04) | M4 (9.99E-04) | M2 (6.99E-03) |
| lead diacetate trihydrate | 3.9$\mu$M | kidney | M4, M7 | **M4** (9.99E-04) | M5 (9.99E-04) | M2 (9.99E-04) |
| lomustine | 4.8$\mu$M | hepatocytes | M5, M6 | M4 (9.99E-04) | M1 (9.99E-04) | **M5** (9.99E-04) |
| lomustine | 24$\mu$M | hepatocytes | M5, M6 | M1 (9.99E-04) | M4 (9.99E-04) | **M5** (9.99E-04) |
| lomustine | 120$\mu$M | hepatocytes | M5, M6 | M4 (9.99E-04) | M2 (9.99E-04) | **M5** (9.99E-04) |
| menthol | 1mM | kidney | M5 | **M5** (9.99E-04) | M4 (9.99E-04) | M2 (9.99E-04) |
| methapyrilene | 24$\mu$M | hepatocytes | M1, M5 | M2 (9.99E-04) | M8 (9.99E-04) | N/A |
| methapyrilene | 120$\mu$M | hepatocytes | M1, M5 | **M1** (9.99E-04) | M4 (9.99E-04) | **M5** (8.99E-03) |
| methapyrilene | 600$\mu$M | hepatocytes | M1, M5 | M4 (9.99E-04) | M2 (9.99E-04) | **M5** (9.99E-04) |
| methyltestosterone | 0.8$\mu$M | hepatocytes | M8 | M4 (9.99E-04) | M1 (9.99E-04) | M5 (9.99E-04) |

| | | | | | | |
|---|---|---|---|---|---|---|
| methyltestosterone | 4$\mu$M | hepatocytes | M8 | M1 (9.99E-04) | M4 (9.99E-04) | M5 (9.99E-04) |
| methyltestosterone | 20$\mu$M | hepatocytes | M8 | M4 (9.99E-04) | M5 (9.99E-04) | M2 (9.99E-04) |
| naphthyl isothiocyanate | 8$\mu$M | hepatocytes | M4, M5 | **M4** (9.99E-04) | M1 (9.99E-04) | **M5** (9.99E-04) |
| naphthyl isothiocyanate | 40$\mu$M | hepatocytes | M4, M5 | **M4** (9.99E-04) | **M5** (9.99E-04) | M1 (9.99E-04) |
| naphthyl isothiocyanate | 200$\mu$M | hepatocytes | M4, M5 | M1 (9.99E-04) | **M4** (9.99E-04) | **M5** (9.99E-04) |
| nitrofurantoin | 5$\mu$M | hepatocytes | M4, M5 | M2 (9.99E-04) | M1 (9.99E-04) | **M5** (1.10E-02) |
| nitrofurantoin | 25$\mu$M | hepatocytes | M4, M5 | M2 (9.99E-04) | **M4** (9.99E-04) | **M5** (9.99E-04) |
| nitrofurantoin | 125$\mu$M | hepatocytes | M4, M5 | **M4** (9.99E-04) | M1 (9.99E-04) | **M5** (9.99E-04) |
| N-methyl-N-nitrosurea | 500$\mu$M | lung epithelial | M2, M3 | M5 (9.99E-04) | M4 (9.99E-04) | M1 (9.99E-04) |
| N-methyl-N-nitrosurea | 1000$\mu$M | lung epithelial | M2, M3 | M5 (9.99E-04) | M4 (9.99E-04) | **M2** (9.99E-04) |
| N-nitrosomorpholine | 183$\mu$M | kidney | M5 | M4 (9.99E-04) | M1 (9.99E-04) | **M5** (2.00E-03) |
| N-nitrosomorpholine | 2500$\mu$M | lung epithelial | M5 | **M5** (9.99E-04) | M4 (9.99E-04) | M2 (9.99E-04) |
| N-nitrosomorpholine | 5000$\mu$M | lung epithelial | M5 | M4 (9.99E-04) | **M5** (9.99E-04) | M2 (9.99E-04) |
| omeprazole | 24$\mu$M | hepatocytes | M4, M5 | **M5** (9.99E-04) | M1 (9.99E-04) | M2 (9.99E-04) |
| omeprazole | 120$\mu$M | hepatocytes | M4, M5 | **M4** (9.99E-04) | M2 (9.99E-04) | **M5** (9.99E-04) |
| omeprazole | 600$\mu$M | hepatocytes | M4, M5 | **M4** (9.99E-04) | M1 (9.99E-04) | **M5** (9.99E-04) |
| perhexiline | 0.6$\mu$M | hepatocytes | M5 | M10 (3.20E-02) | N/A | N/A |
| perhexiline | 3$\mu$M | hepatocytes | M5 | M1 (5.00E-03) | M3 (9.99E-04) | M4 (2.00E-03) |
| perhexiline | 15$\mu$M | hepatocytes | M5 | M4 (9.99E-04) | **M5** (9.99E-04) | M2 (9.99E-04) |
| phenobarbital | 400$\mu$M | hepatocytes | M4 | **M4** (9.99E-04) | M1 (9.99E-04) | M5 (9.99E-04) |
| phenobarbital | 2000$\mu$M | hepatocytes | M4 | **M4** (9.99E-04) | M5 (9.99E-04) | M2 (9.99E-04) |
| phenobarbital | 10000$\mu$M | hepatocytes | M4 | M5 (9.99E-04) | **M4** (9.99E-04) | M2 (9.99E-04) |
| phenobarbital sodium | 780$\mu$M | HepG2 | M4 | M3 (9.99E-04) | M8 (2.00E-03) | M6 (2.90E-02) |
| phenobarbital sodium | 1560$\mu$M | HepG2 | M4 | M1 (2.00E-03) | M2 (3.00E-02) | M8 (9.99E-04) |
| phenylbutazone | 16$\mu$M | hepatocytes | M4, M5 | M2 (9.99E-04) | **M5** (7.99E-03) | M8 (3.20E-02) |
| phenylbutazone | 80$\mu$M | hepatocytes | M4, M5 | **M4** (9.99E-04) | **M5** (9.99E-04) | M2 (9.99E-04) |
| phenylbutazone | 400$\mu$M | hepatocytes | M4, M5 | **M4** (9.99E-04) | **M5** (9.99E-04) | M2 (9.99E-04) |

| | | | | | | |
|---|---|---|---|---|---|---|
| phenytoin | 2.4$\mu$M | hepatocytes | M4 | M2 (9.99E-04) | M8 (9.99E-04) | N/A |
| phenytoin | 12$\mu$M | hepatocytes | M4 | M1 (9.99E-04) | M5 (4.70E-02) | M2 (9.99E-04) |
| phenytoin | 60$\mu$M | hepatocytes | M4 | **M4** (9.99E-04) | M2 (9.99E-04) | M1 (9.99E-04) |
| phorbol 13-acetate 12-myristate | 1E07$\mu$M | HepG2 | M8 | M4 (9.99E-04) | M1 (9.99E-04) | M2 (9.99E-04) |
| phorbol 13-acetate 12-myristate | 35$\mu$M | HepaRG | M8 | M4 (9.99E-04) | M5 (9.99E-04) | M1 (9.99E-04) |
| piperonyl butoxide | 3.2$\mu$M | HepaRG | M4 | M11 (4.40E-02) | N/A | N/A |
| piperonyl butoxide | 22.5$\mu$M | HepG2 | M4 | M5 (9.99E-04) | N/A | N/A |
| piperonyl butoxide | 45$\mu$M | HepG2 | M4 | M5 (9.99E-04) | N/A | N/A |
| pirinixic acid (WY-14643) | 0.09$\mu$M | HepG2 | M2, M5 | M4 (9.99E-04) | **M5** (9.99E-04) | **M2** (9.99E-04) |
| pirinixic acid (WY-14643) | 6$\mu$M | hepatocytes | M2, M5 | M4 (9.99E-04) | **M5** (9.99E-04) | **M2** (9.99E-04) |
| pirinixic acid (WY-14643) | 30$\mu$M | hepatocytes | M2, M5 | M4 (9.99E-04) | **M2** (9.99E-04) | **M5** (9.99E-04) |
| pirinixic acid (WY-14643) | 150$\mu$M | hepatocytes | M2, M5 | M4 (9.99E-04) | M1 (9.99E-04) | **M5** (9.99E-04) |
| pirinixic acid (WY-14643) | 210$\mu$M | HepaRG | M2, M5 | M4 (9.99E-04) | M1 (9.99E-04) | **M5** (1.20E-02) |
| potassium bromate | 1mM | kidney | M4 | **M4** (9.99E-04) | M5 (9.99E-04) | M2 (9.99E-04) |
| propylthiouracil | 160$\mu$M | hepatocytes | M4 | **M4** (9.99E-04) | M5 (9.99E-04) | M2 (9.99E-04) |
| propylthiouracil | 800$\mu$M | hepatocytes | M4 | **M4** (9.99E-04) | M5 (9.99E-04) | M2 (9.99E-04) |
| propylthiouracil | 4000$\mu$M | hepatocytes | M4 | **M4** (9.99E-04) | M1 (9.99E-04) | M5 (9.99E-04) |
| resorcinol | 500$\mu$M | lung epithelial | M5 | M4 (9.99E-04) | **M5** (9.99E-04) | M2 (9.99E-04) |
| resorcinol | 1000$\mu$M | lung epithelial | M5 | M4 (9.99E-04) | M2 (9.99E-04) | M1 (9.99E-04) |
| rifampicin | 2.8$\mu$M | hepatocytes | M4, M11 | M1 (9.99E-04) | **M4** (9.99E-04) | M5 (9.99E-04) |
| rifampicin | 14$\mu$M | hepatocytes | M4, M11 | M1 (9.99E-04) | **M4** (9.99E-04) | M5 (9.99E-04) |
| rifampicin | 70$\mu$M | hepatocytes | M4, M11 | **M4** (9.99E-04) | M5 (9.99E-04) | M2 (9.99E-04) |
| rotenone | 2$\mu$M | hepatocytes | M5 | **M5** (9.99E-04) | M1 (9.99E-04) | M9 (9.99E-04) |
| sodium dichromate | 2.5$\mu$M | lung epithelial | M4 | M5 (9.99E-04) | **M4** (9.99E-04) | M1 (9.99E-04) |
| sodium dichromate | 5$\mu$M | lung epithelial | M4 | **M4** (9.99E-04) | M5 (9.99E-04) | M2 (9.99E-04) |

| | | | | | | |
|---|---|---|---|---|---|---|
| sodium metaarsenite | 3.75$\mu$M | lung epithelial | M4 | **M4** (9.99E-04) | M5 (9.99E-04) | M1 (9.99E-04) |
| sodium metaarsenite | 7.5$\mu$M | lung epithelial | M4 | **M4** (9.99E-04) | M1 (9.99E-04) | M5 (9.99E-04) |
| streptozocin | 10000$\mu$M | kidney | M1, M5 | M4 (9.99E-04) | M2 (9.99E-04) | **M5** (9.99E-04) |
| styrene | 3000$\mu$M | lung epithelial | M4 | **M4** (9.99E-04) | M2 (9.99E-04) | M1 (9.99E-04) |
| styrene | 6000$\mu$M | lung epithelial | M4 | **M4** (9.99E-04) | M5 (9.99E-04) | M2 (9.99E-04) |
| sulfasalazine | 6$\mu$M | hepatocytes | M4 | M5 (9.99E-04) | M2 (9.99E-04) | M1 (6.99E-03) |
| sulfasalazine | 30$\mu$M | hepatocytes | M4 | M5 (4.30E-02) | M2 (1.20E-02) | M3 (9.99E-04) |
| sulfasalazine | 150$\mu$M | hepatocytes | M4 | **M4** (9.99E-04) | M5 (9.99E-04) | M2 (9.99E-04) |
| S-(1,2-dichlorovinyl)-L-cysteine | 60$\mu$M | kidney | M5 | **M5** (9.99E-04) | M1 (9.99E-04) | M8 (9.99E-04) |
| tacrine | 80$\mu$M | hepatocytes | M5 | M1 (9.99E-04) | **M5** (2.00E-03) | M2 (9.99E-04) |
| tetracycline | 1$\mu$M | hepatocytes | M5 | M2 (9.99E-04) | **M5** (9.99E-04) | M1 (9.99E-04) |
| tetracycline | 5$\mu$M | hepatocytes | M5 | M10 (2.40E-02) | N/A | N/A |
| tetracycline | 25$\mu$M | hepatocytes | M5 | M2 (9.99E-04) | **M5** (9.99E-04) | M1 (9.99E-04) |
| thioacetamide | 400$\mu$M | hepatocytes | M4, M5 | **M4** (9.99E-04) | **M5** (9.99E-04) | M2 (9.99E-04) |
| thioacetamide | 2000$\mu$M | hepatocytes | M4, M5 | **M5** (9.99E-04) | M1 (9.99E-04) | M2 (9.99E-04) |
| thioacetamide | 10000$\mu$M | hepatocytes | M4, M5 | **M4** (9.99E-04) | M1 (9.99E-04) | **M5** (9.99E-04) |
| thioridazine | 0.6$\mu$M | hepatocytes | M1, M6, M7 | N/A | N/A | N/A |
| thioridazine | 3$\mu$M | hepatocytes | M1, M6, M7 | **M1** (5.00E-03) | M4 (9.99E-04) | M2 (9.99E-04) |
| thioridazine | 15$\mu$M | hepatocytes | M1, M6, M7 | M4 (9.99E-04) | **M1** (9.99E-04) | M5 (4.00E-03) |
| tolbutamide | 2.1$\mu$M | HepG2 | M2 | M4 (9.99E-04) | **M2** (9.99E-04) | M1 (9.99E-04) |
| tolbutamide | 20$\mu$M | kidney | M2 | M1 (9.99E-04) | M4 (9.99E-04) | **M2** (9.99E-04) |
| tolbutamide | 2000$\mu$M | HepaRG | M2 | M4 (9.99E-04) | M5 (9.99E-04) | **M2** (9.99E-04) |
| triclosan | 250nM | kidney | M1, M8 | M2 (9.99E-04) | M4 (9.99E-04) | **M1** (9.99E-04) |
| tris(2,3-dibromopropyl)phosphate | 1.26$\mu$M | kidney | M4 | **M4** (9.99E-04) | M5 (9.99E-04) | M1 (4.00E-03) |
| valproic acid | 200$\mu$M | hepatocytes | M4, M5 | **M4** (9.99E-04) | **M5** (9.99E-04) | M1 (9.99E-04) |

| | | | | | | |
|---|---|---|---|---|---|---|
| valproic acid | 1000$\mu$M | hepatocytes | M4, M5 | **M4** (9.99E-04) | M2 (9.99E-04) | **M5** (9.99E-04) |
| valproic acid | 5000$\mu$M | hepatocytes | M4, M5 | **M4** (9.99E-04) | M1 (9.99E-04) | **M5** (9.99E-04) |

These are the three toxicity mechanisms with the highest enrichment scores, for each of the public gene expression (transcriptomics) assays that were tested. It is worth noting that several of these assays may not be conducted at a toxic dose, and thus may result either in an erroneous prediction or no predictions at all, due to a lack of significantly enriched mechanisms. The **bolded** predictions are those that match the known mechanisms for that chemical.
