## Supplementary material for "Applying knowledge-driven mechanistic inference to toxicogenomics": supplemental_table_S3.pdf

**MechSpy predictions for chemicals for which there is no strong enough evidence of a particular mechanism of toxicity.**

| Chemical | Dose | Cell Type | #1 Predicted mechanism | #2 Predicted mechanism | #3 Predicted mechanism |
|---|---|---|---|---|---|
| 1-Amino-2,4-dibromoanthra-quinone | 150$\mu$M | kidney | M4 (9.99E-04) | M5 (9.99E-04) | M2 (9.99E-04) |
| 2-Amino-3-methylimidazo(4,5-f)quinoline | 2.9$\mu$M | HepaRG | M4 (9.99E-04) | M5 (9.99E-04) | M1 (9.99E-04) |
| 2-nitrofluorene | 9$\mu$M | HepaRG | M2 (1.10E-02) | M3 (2.60E-02) | N/A |
| 2-nitrofluorene | 18$\mu$M | HepaRG | M4 (9.99E-04) | M1 (9.99E-04) | M5 (8.99E-03) |
| 2-nitrofluorene | 35$\mu$M | HepG2 | N/A | N/A | N/A |
| 2-nitrofluorene | 60$\mu$M | kidney | N/A | N/A | N/A |
| 4-Acetylaminofluorene | 220$\mu$M | kidney | M4 (9.99E-04) | M2 (9.99E-04) | M5 (9.99E-04) |
| 4-Acetylaminofluorene | 222.2$\mu$M | HepaRG | M5 (7.99E-03) | N/A | N/A |
| adapin | 3$\mu$M | hepatocytes | M2 (9.99E-04) | M8 (2.00E-03) | N/A |
| adapin | 15$\mu$M | hepatocytes | M1 (9.99E-04) | M5 (2.00E-03) | M4 (9.99E-04) |
| adapin | 75$\mu$M | hepatocytes | M4 (9.99E-04) | M1 (9.99E-04) | M5 (9.99E-04) |
| aspirin | 120$\mu$M | hepatocytes | N/A | N/A | N/A |
| beclomethasone dipropionate | 50$\mu$M | lung epithelial | M4 (9.99E-04) | M5 (9.99E-04) | M1 (9.99E-04) |
| beclomethasone dipropionate | 100$\mu$M | lung epithelial | M5 (9.99E-04) | M4 (9.99E-04) | M1 (9.99E-04) |
| benzofuran | 250$\mu$M | lung epithelial | M5 (9.99E-04) | M4 (9.99E-04) | M2 (9.99E-04) |
| benzofuran | 500$\mu$M | lung epithelial | M4 (9.99E-04) | M2 (9.99E-04) | M1 (9.99E-04) |
| benzoin | 470$\mu$M | kidney | M4 (9.99E-04) | M1 (9.99E-04) | M5 (9.99E-04) |

| | | | | | |
|---|---|---|---|---|---|
| bromodichloromethane | 200$\mu$M | kidney | M2 (9.99E-04) | M4 (9.99E-04) | M9 (9.99E-04) |
| chlorpromazine | 0.8$\mu$M | hepatocytes | M5 (9.99E-04) | M1 (6.99E-03) | M2 (2.10E-02) |
| chlorpromazine | 4$\mu$M | hepatocytes | M4 (9.99E-04) | M5 (2.00E-03) | M2 (9.99E-04) |
| chlorpromazine | 20$\mu$M | hepatocytes | M4 (9.99E-04) | M1 (9.99E-04) | M5 (9.99E-04) |
| cimetidine | 60$\mu$M | hepatocytes | M8 (3.50E-02) | N/A | N/A |
| cimetidine | 300$\mu$M | hepatocytes | M4 (9.99E-04) | M2 (9.99E-04) | M1 (4.00E-03) |
| dimethyl sulfoxide | 0.0050% | lung epithelial | M4 (9.99E-04) | M5 (9.99E-04) | M2 (9.99E-04) |
| ethionine | 400$\mu$M | hepatocytes | M4 (9.99E-04) | M1 (9.99E-04) | M5 (9.99E-04) |
| ethionine | 2000$\mu$M | hepatocytes | M4 (9.99E-04) | M1 (9.99E-04) | M5 (9.99E-04) |
| ethionine | 10000$\mu$M | hepatocytes | M4 (9.99E-04) | M5 (9.99E-04) | M2 (9.99E-04) |
| hydrazine dihydrochloride | 0.85mM | HepaRG | M4 (9.99E-04) | M5 (9.99E-04) | M1 (9.99E-04) |
| hydroxyzine | 6$\mu$M | hepatocytes | N/A | N/A | N/A |
| hydroxyzine | 30$\mu$M | hepatocytes | M8 (9.99E-04) | N/A | N/A |
| hydroxyzine | 150$\mu$M | hepatocytes | M9 (9.99E-04) | M8 (9.99E-04) | N/A |
| interleukin-6,-human | 2$\mu$M | hepatocytes | M4 (9.99E-04) | M1 (9.99E-04) | M5 (9.99E-04) |
| interleukin-6,-human | 10$\mu$M | hepatocytes | M4 (9.99E-04) | M1 (9.99E-04) | M5 (9.99E-04) |
| interleukin-6,-human | 50$\mu$M | hepatocytes | M4 (9.99E-04) | M5 (9.99E-04) | M2 (9.99E-04) |
| ipratropium bromide hydrate | 625$\mu$M | lung epithelial | M4 (9.99E-04) | M2 (9.99E-04) | M1 (9.99E-04) |
| ipratropium bromide hydrate | 1250$\mu$M | lung epithelial | M4 (9.99E-04) | M5 (9.99E-04) | M1 (9.99E-04) |
| labetalol | 5.6$\mu$M | hepatocytes | M2 (5.00E-03) | M8 (4.00E-03) | N/A |
| labetalol | 28$\mu$M | hepatocytes | M4 (9.99E-04) | M1 (6.99E-03) | M5 (1.80E-02) |
| labetalol | 140$\mu$M | hepatocytes | M1 (9.99E-04) | M4 (9.99E-04) | M2 (1.10E-02) |
| methapyrilene hydrochloride | 75$\mu$M | HepaRG | N/A | N/A | N/A |
| monuron | 200$\mu$M | kidney | N/A | N/A | N/A |
| nifedipine | 20$\mu$M | HepaRG | N/A | N/A | N/A |
| nifedipine | 40$\mu$M | HepaRG | N/A | N/A | N/A |
| nifedipine | 210$\mu$M | HepG2 | M5 (9.99E-04) | M4 (9.99E-04) | M2 (9.99E-04) |

| | | | | | |
|---|---|---|---|---|---|
| nifedipine | 10$\mu$M | kidney | M4 (9.99E-04) | M5 (9.99E-04) | M2 (9.99E-04) |
| nitrilotriacetic-acid | 260$\mu$M | kidney | M2 (2.60E-02) | M9 (9.99E-04) | M10 (2.00E-03) |
| N-Ethyl-N-(2-hydroxyethyl)nitrosamine | 8.45$\mu$M | kidney | M4 (9.99E-04) | M1 (9.99E-04) | M5 (2.00E-03) |
| ochratoxin-A | 300nM | kidney | M4 (9.99E-04) | M5 (9.99E-04) | M2 (9.99E-04) |
| phthalic anhydride | 2000$\mu$M | lung epithelial | M4 (9.99E-04) | M5 (9.99E-04) | M2 (9.99E-04) |
| phthalic anhydride | 4000$\mu$M | lung epithelial | M4 (9.99E-04) | M5 (9.99E-04) | M2 (9.99E-04) |
| THS 2.2 | 0.15mg/l | nasal epithelial | M5 (9.99E-04) | M8 (9.99E-04) | M10 (2.40E-02) |
| THS 2.2 | 0.27mg/l | nasal epithelial | M10 (9.99E-04) | N/A | N/A |
| THS 2.2 | 0.31mg/l | buccal epithelial | M1 (1.80E-02) | M8 (9.99E-04) | M7 (9.99E-04) |
| THS 2.2 | 0.44mg/l | nasal epithelial | M8 (4.00E-03) | N/A | N/A |
| THS 2.2 | 0.46mg/l | buccal epithelial | M1 (2.80E-02) | M10 (9.99E-04) | N/A |
| THS 2.2 | 1.09mg/l | buccal epithelial | M4 (9.99E-04) | M5 (9.99E-04) | M1 (9.99E-04) |
| THS 2.2 | 7.5$\mu$g/l | bronchial epithelial | M1 (9.99E-04) | M4 (9.99E-04) | M5 (9.99E-04) |
| THS 2.2 | 37.5$\mu$g/l | bronchial epithelial | M4 (9.99E-04) | M5 (9.99E-04) | M1 (9.99E-04) |
| THS 2.2 | 150$\mu$g/l | bronchial epithelial | M4 (9.99E-04) | M5 (9.99E-04) | M1 (9.99E-04) |
| TGF$\beta$1 | 2$\mu$M | hepatocytes | M4 (9.99E-04) | M1 (9.99E-04) | M5 (9.99E-04) |
| TGF$\beta$1 | 10$\mu$M | hepatocytes | M1 (9.99E-04) | M4 (9.99E-04) | M5 (9.99E-04) |
| TGF$\beta$1 | 50$\mu$M | hepatocytes | M4 (9.99E-04) | M1 (9.99E-04) | M5 (9.99E-04) |
| urea | 5000$\mu$M | lung epithelial | M4 (9.99E-04) | M2 (9.99E-04) | M1 (9.99E-04) |
| urea | 10000$\mu$M | lung epithelial | M4 (9.99E-04) | M5 (9.99E-04) | M2 (9.99E-04) |

Some of MechSpy-generated hypotheses were experimentally validated for two of these chemicals (adapin and chlorpromazine).
